# Climate warming suppresses soil abundant fungal taxa and reduces soil carbon efflux in a semi-arid grassland

**DOI:** 10.1101/2023.10.08.561373

**Authors:** Yunpeng Qiu, Kangcheng Zhang, Yunfeng Zhao, Yexin Zhao, Bianbian Wang, Yi Wang, Tangqing He, Xinyu Xu, Tongshuo Bai, Yi Zhang, Shuijin Hu

**Author notes:** These authors contributed equally to this work (Yunpeng Qiu, Kangcheng Zhang, Yunfeng Zhao). Corresponding author: Yunpeng Qiu, Shuijin Hu.

## Abstract

The abundance, diversity and activity of soil microorganisms critically control the fate of recent plant carbon (C) inputs as well as protecting soil organic C, regulating C-climate feedbacks. However, the effects of climate change drivers such as warming and precipitation change on soil microbial communities and C dynamics remain poorly understood. Utilizing a long-term field warming and precipitation manipulation in a semi-arid grassland on the Loess Plateau and a complementary incubation experiment, here we show that warming and rainfall reduction differentially affect the abundance and composition of bacteria and fungi, and soil C efflux. Warming significantly reduced the abundance of fungi but not bacteria, increasing the relative dominance of bacteria in the soil microbial community. In particular, warming shifted the community composition of abundant fungi in favor of oligotrophic *Capnodiales* and *Hypocreales* over potential saprotroph *Archaeorhizomycetales*. In contrast, precipitation reduction increased soil microbial biomass, but did not significantly affect either the abundance or diversity of bacteria. Furthermore, soil abundant, not rare, fungal community composition was significantly correlated with soil CO_2_ efflux. Our findings suggest that alterations in the fungal community composition, in response to changes in soil C and moisture, dominate the microbial responses to climate change and thus control soil C dynamics in semi-arid grasslands.

**Impact statement:** Semi-arid grasslands play a critical role in global carbon (C) cycle and potential feedbacks to climate change. Understanding the responses of soil microorganisms to warming and rainfall change is key to evaluating and predicting semi-arid grassland soil C dynamics under future climate change scenarios. Our study showed that warming induced a shift in the abundant fungal community, favoring oligotrophic fungi (i.e., *Capnodiales* and *Hypocreales*) over the potential saprotrophic *Archaeorhizomycetales,* and reduced C efflux. These findings advance our understanding of soil microbial and C responses to climate change drivers and may help predict and possibly manage soil C sequestration in semi-arid grasslands.

## INTRODUCTION

Soil microorganisms play a vital role in carbon (C) cycling through controlling the fate of recent plant C inputs as well as protecting soil organic carbon, while mineralizing nutrients to sustain plant C fixation^1,2^. Soil bacteria and fungi constitute the most diverse and abundant microbial communities on earth and dominate the litter decomposition and nutrient cycling^3,4^. Soil bacteria and fungi have distinct morphologies, cell structures and physiological traits, and respond divergently to warming and precipitation reduction across different ecosystems^1,5,6^. Compared with bacteria, fungi are more resistant to water stress induced by climate warming and reduced precipitation because they have thick cell wall and extensive hyphal networks to access nutrient and water over long distances^7,8^. Similarly, Gram-positive (G^+^) bacteria with strong, thick and interlinked peptidoglycan cell walls are inherently resistant to warm and dry conditions^7,9^. Physiologically, bacteria and fungi have different requirements for C and nitrogen (N), and C use efficiency, as fungal biomass have a higher C:N ratio, and high fungal to bacteria abundances are believed to be conducive for soil C retention^3,10^.

Bacterial and fungal communities are highly diverse with both abundant and rare taxa that are often with the unbalanced distribution of large numbers of rare species versus a few highly abundant species^11,12^. Both abundant and rare microbial taxa contribute to degrading and mineralizing organic materials and thus play a critical role in regulating the C-climate feedbacks^13–15^. Previous studies have largely been focused on the abundant members of the soil microbial communities because they often dominate the microbial biomass and are presumably play the dominant role in biogeochemical C and nutrient cycling^14,16^. Emerging evidence has, however, shown that rare species may play a disproportionately important role in a change climate^13,15^. Despite its low abundance, rare taxa provide a tremendous reservoir of genetic and functional diversity that allows microbial communities to respond rapidly to environmental change^13,17,18^, and to maintain ecosystem stability and function^19,20^. For example, Xiong et al. (2014)^17^ found that rare species of soil fungi are particularly sensitive to warming in an alpine meadow on Tibetan Plateau. Similarly, precipitation reduction can also alter soil abundant and rare microbial taxa differentially^8,21^. In a subtropical forest, Zhao et al. (2017)^21^ reported that precipitation reduction impacted the dominant fungal but rare bacterial taxa. Meanwhile, rare members are likely to be functionally dissimilar from the abundant members, which may enhance functionality of the abundant microorganisms^13,22^. Therefore, understanding the differential responses of abundant and rare taxa to climate change drivers may be key to better predicting terrestrial C dynamics and the carbon-climate feedbacks under future climate change scenarios.

Global surface temperature has increased by around 1.09 °C since the industrialization and is projected to continue to increase by another 1.4-5.8°C by the end of this century^23^. Climate warming is often associated with changes in water balance through enhancing evapotranspiration and altering precipitation regimes^23,24^. Among the global regions, most vulnerable to the ongoing climate change are those at the mid- to high latitude, such as the Loess Plateau in Northwest China^25,26^. The Loess Plateau covers an area of ca. 640,000 km^2^ and its deep soil largely originates from the accumulation of wind-blown dust^27^. In addition of long-term human disturbance, low rainfall with extreme uneven distribution limits plant growth, leading to the most serious soil erosion in the world^28^. Over the last five decades, the average temperature on the Loess Plateau has increased by ca. 1.9 °C^25^, as compared to the increase of 1.09 °C in the average global temperature. Concurrent with the climate warming, this region has also experienced extensive periods of drought followed by heavy precipitation events^26^. Climate warming and precipitation change may alter soil physical and chemical properties (i.e., water content and nutrient substrate), plant growth and plant-derived C input to soil^29,30^. Alterations in soil resources, particularly organic C and soil moisture, can profoundly influence growth and activities of soil microorganisms, and their interactions with the environment, which largely control the fate of recent plant C inputs as well as protecting soil organic C^1,31,32^.

Although many field studies have examined the impact of climate change drivers such as warming and rainfall change on plant productivity and community composition^33,34^, the response of soil microorganisms, the subcommunities of abundant and rare microbes in particular, have received limited attention^18^. We initiated a field experiment in a semi-arid grassland on Loess Plateau to examine the impact of warming, precipitation reduction and their combination on bacterial and fungal communities. Because of severe water limitation with potential high evapotranspiration in the semi-arid ecosystems, warming and precipitation reduction may induce water stress on soil microbial growth. Our objectives were to 1) determine the direct effects of warming and precipitation reduction on soil microbial biomass, activities, and communities of abundant and rare taxa, and 2) examine the potential linkages among plant growth, soil microbial communities, particularly abundant and rare microbial taxa, and soil C dynamics under warming and precipitation reduction. Our hypotheses were that (1) both warming and precipitation reduction are more suppressive to bacteria than fungi, increasing fungal dominance but reducing total microbial CO_2_ efflux, (2) warming and precipitation reduction alter the relative composition of abundant and rare bacteria and fungi in favor of rare taxa, and (3) alterations in the microbial community composition correlate with soil CO_2_.

## RESULTS

### Responses of soil chemical, microbial properties and C efflux to warming and precipitation reduction

Across all the treatments, NO_3-_-N (11.1 mg N kg^-1^ soil) was more dominant than NH_4+_-N (2.41 mg N kg^-1^ soil) (Table 1). Warming, but not precipitation reduction, significantly reduced soil NH^+^-N by 16.2% (Table 1). Warming alone tended to increase soil NO_3-_-N by 16.2% and precipitation reduction significantly enhanced soil NO_3-_-N (Table 1). While precipitation reduction had no effect on soil dissolved organic carbon (DOC), net mineralization rate (NMR) or microbial biomass carbon (MBC), warming significantly enhanced soil DOC but reduced MBC and NMR (all *p* < 0.05; Table 1). Also, warming significantly reduced the MBC:MBN ratio (*p* < 0.01; Table 1), suggesting a decrease in the relative dominance of fungi over bacteria. In contrast, precipitation reduction significantly reduced MBN, leading to a marginal increase in the ratio of MBC to MBN (*p* = 0.06; Table 1).

**Table 1.** Effects of warming and precipitation reduction on soil properties. Data presented as mean values with standard errors (n = 4).

| Variables | Treatments |  |  |  | Two-way ANOVA |  |  |
| --- | --- | --- | --- | --- | --- | --- | --- |
|  | CK | Pr | W | WPr | W | PR | W×Pr |
| NH<sub>4</sub><sup>+</sup>-N (mg N kg<sup>-1</sup>) | 2.57 ± 0.24 | 2.67 ± 0.26 | 2.49 ± 0.03 | 1.90 ± 0.08 | **<0.05** | 0.21 | *0.08* |
| NO<sub>3</sub><sup>-</sup>-N (mg N kg<sup>-1</sup>) | 9.0 ± 0.5 | 12.9 ± 1.4 | 10.5 ± 0.6 | 12.1 ± 1.1 | 0.72 | **<0.05** | 0.26 |
| DOC (mg C kg<sup>-1</sup>) | 140.4 ± 3.4 | 151.8 ± 20.2 | 173.4 ± 9.5 | 176.7 ± 10.5 | **<0.05** | 0.57 | 0.75 |
| NMR (mg N kg<sup>-1</sup> d<sup>-1</sup>) | 1.19 ± 0.14 | 1.18 ± 0.22 | 0.84 ± 0.04 | 0.90 ± 0.06 | **<0.05** | 0.85 | 0.80 |
| MBC (mg C kg<sup>-1</sup>) | 698.2 ± 67.6 | 803.5 ± 40.0 | 584.0 ± 79.7 | 511.2 ± 89.8 | **<0.05** | 0.83 | 0.24 |
| MBN (mg N kg<sup>-1</sup>) | 160.4 ± 20.9 | 118.2 ± 3.9 | 159.1 ± 8.2 | 125.5±14.4 | 0.83 | **<0.05** | 0.75 |
| MBC/MBN | 4.7 ± 0.9 | 6.8 ± 0.2 | 3.7 ± 0.5 | 4.0 ± 0.6 | **<0.01** | *0.06* | 0.16 |

DOC, dissolved organic carbon; NMR, soil net nitrogen mineralization rates; MBC, soil microbial biomass carbon; MBN, soil microbial biomass nitrogen. CK, control; W, warming; Pr, precipitation reduction; WPr, warming + precipitation reduction. Significant effects ( $P < 0.05$ ) are shown in bold, and marginal significant effects ( $0.05 < P \leq 0.10$ ) are in italic, ANOVA mixed model.

We measured soil C efflux through an incubation experiment, using soil samples collected after 5-year field treatments. Soil CO_2_ fluxes varied from 0.165 to 0.365 mg C kg^-1^ soil h^-1^ during the 28-day incubation period and peaked in all the treatments at day 4 (Fig. 1A). On average, warming significantly reduced soil CO_2_ emission by 15.6%, but precipitation reduction had no effect on it (Fig. 1B). There was no significant interaction between warming and precipitation reduction on soil CO_2_ emission (Fig. 1B).

**Fig. 1.**
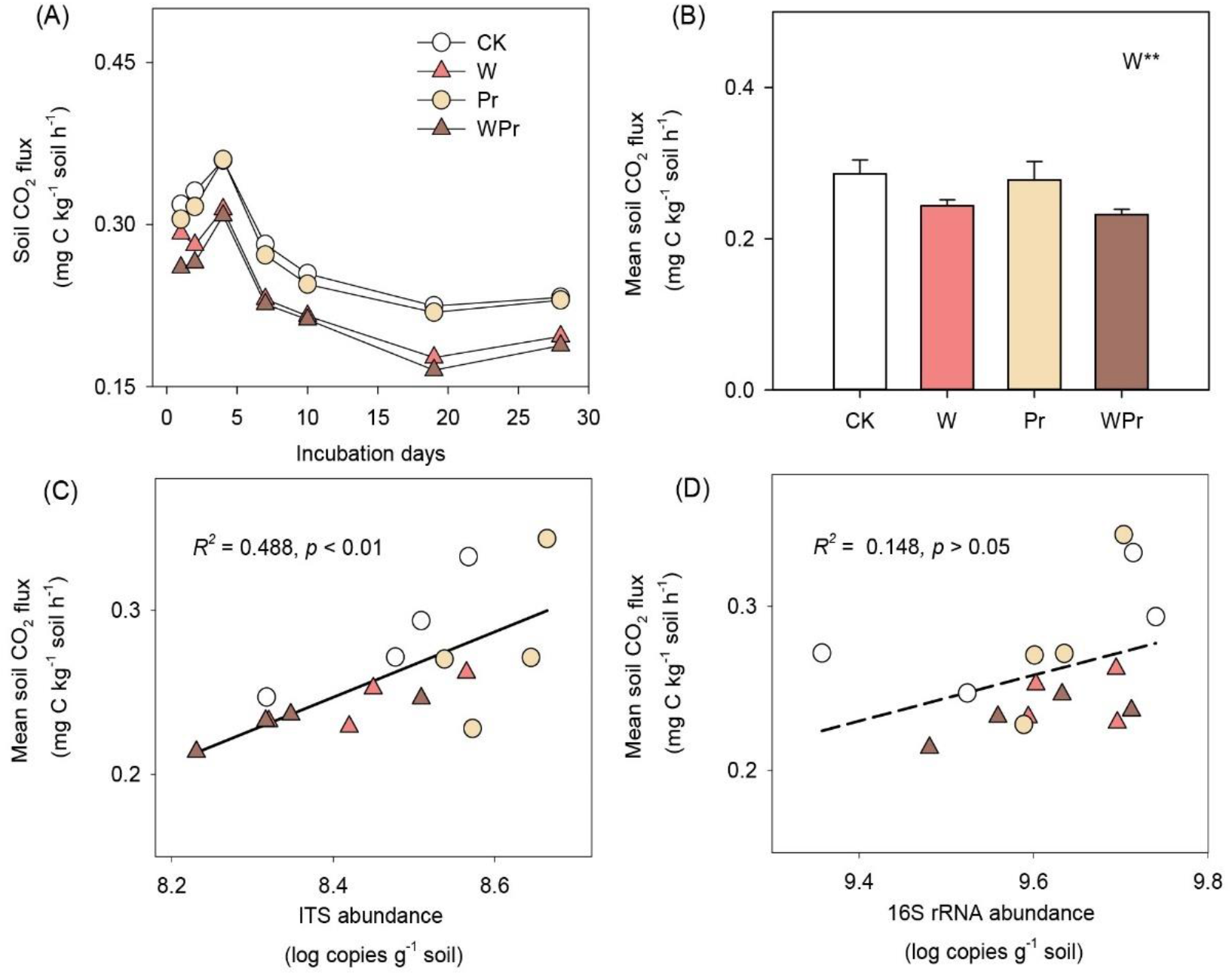
Soil CO_2_ flux and relationship between soil CO_2_ flux and fungal or bacterial abundance. Effects of warming and precipitation reduction on soil CO_2_ flux (a, b), and the relationships between mean soil CO_2_ flux and fungal (c), bacterial abundance (d), respectively. Bars are standard errors of the means (n = 4). The four treatments are as follows: CK, control; W, warming; Pr, precipitation reduction; WPr, combination of warming and precipitation reduction. The significance levels are labeled with **0.001 < *p* ≤ 0.01, ANOVA mixed model.

### Responses of soil bacterial and fungal abundances to warming and precipitation reduction

Neither warming nor precipitation reduction significantly affected soil bacterial abundance (Fig. 2B and Table S2). Warming reduced soil fungal abundance by 27.6% (*p* < 0.05; Fig. 2A and Table S2). There was an interaction between warming and precipitation reduction on soil fungal abundance (*p* < 0.05; Fig. 2A and Table S2). In addition, warming significantly decreased the ratio of fungi to bacteria by 31.1%, although precipitation reduction had no effect on it (Fig. 2C and Table S2)

**Fig. 2.**
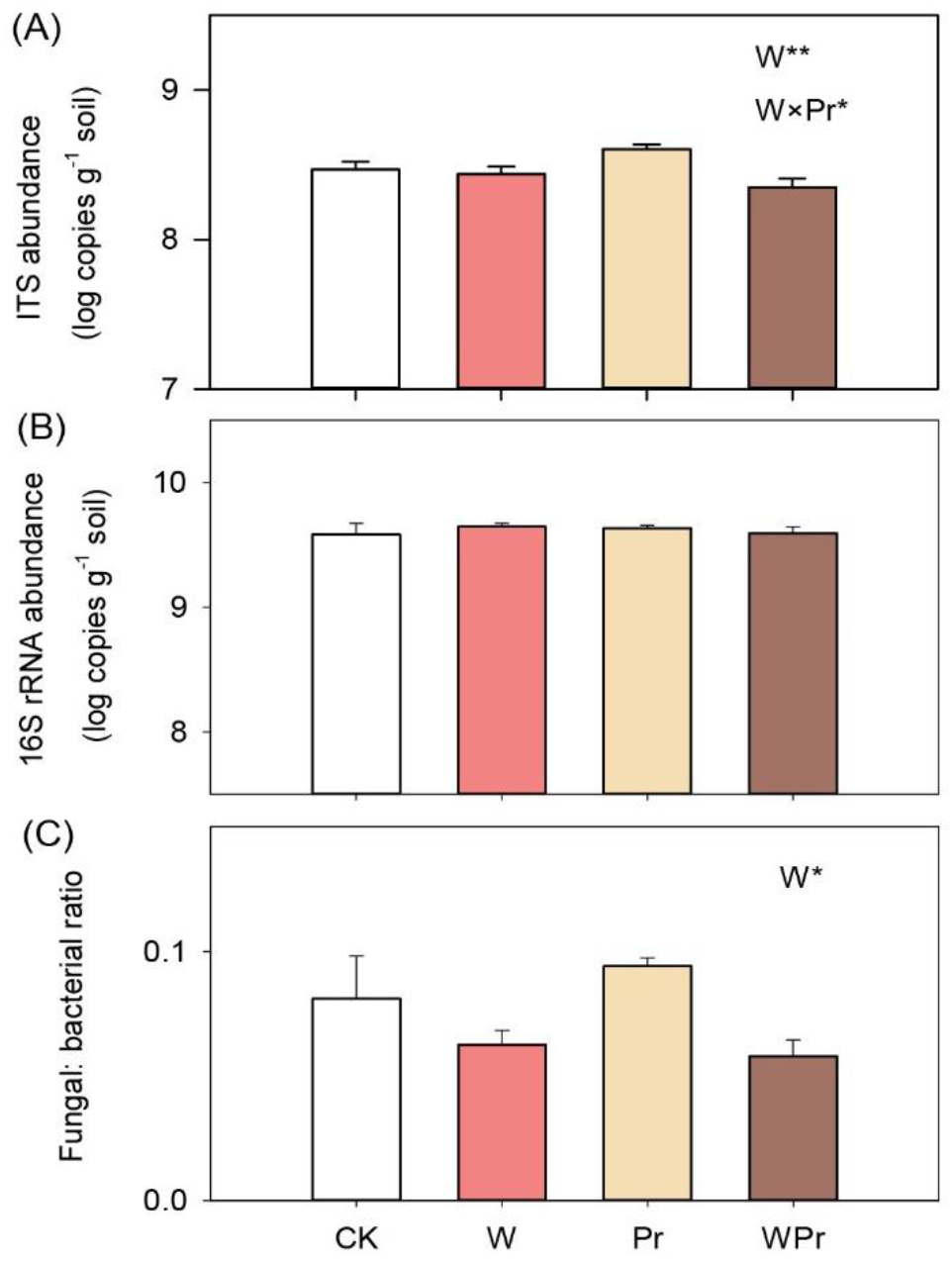
Soil fungal, bacterial abundance and ratio of fungal to bacterial abundance. Effects of warming and precipitation reduction on the abundance of soil bacteria (a), fungi (b) and the ratio of fungi to bacteria (c). Bars are standard errors of the means (n = 4). The four treatments are as follows: CK, control; W, warming; Pr, precipitation reduction; WPr, combination of warming and precipitation reduction. The significance levels are labeled with *0.01 < *p* ≤ 0.05; **0.001 < *p* ≤ 0.01, ANOVA mixed model.

### Responses of abundant and rare soil microbial taxa to warming and precipitation reduction

For soil fungi, while abundant taxa accounted for 77.1% of the sequence reads with a significantly lower proportion of ASVs (8.5%), rare taxa comprised 4.8% of the sequence reads with a significantly higher proportion of ASVs (56.0%) (Fig. 3A, 3B). As for soil bacteria, while abundant taxa accounted for 39.0% of the sequence reads with a significantly lower proportion of ASVs (4.6%), rare taxa comprised 11.2% of the sequence reads with a significantly higher proportion of ASVs (57.2%) (Fig. 3A, 3B).

**Fig. 3.**
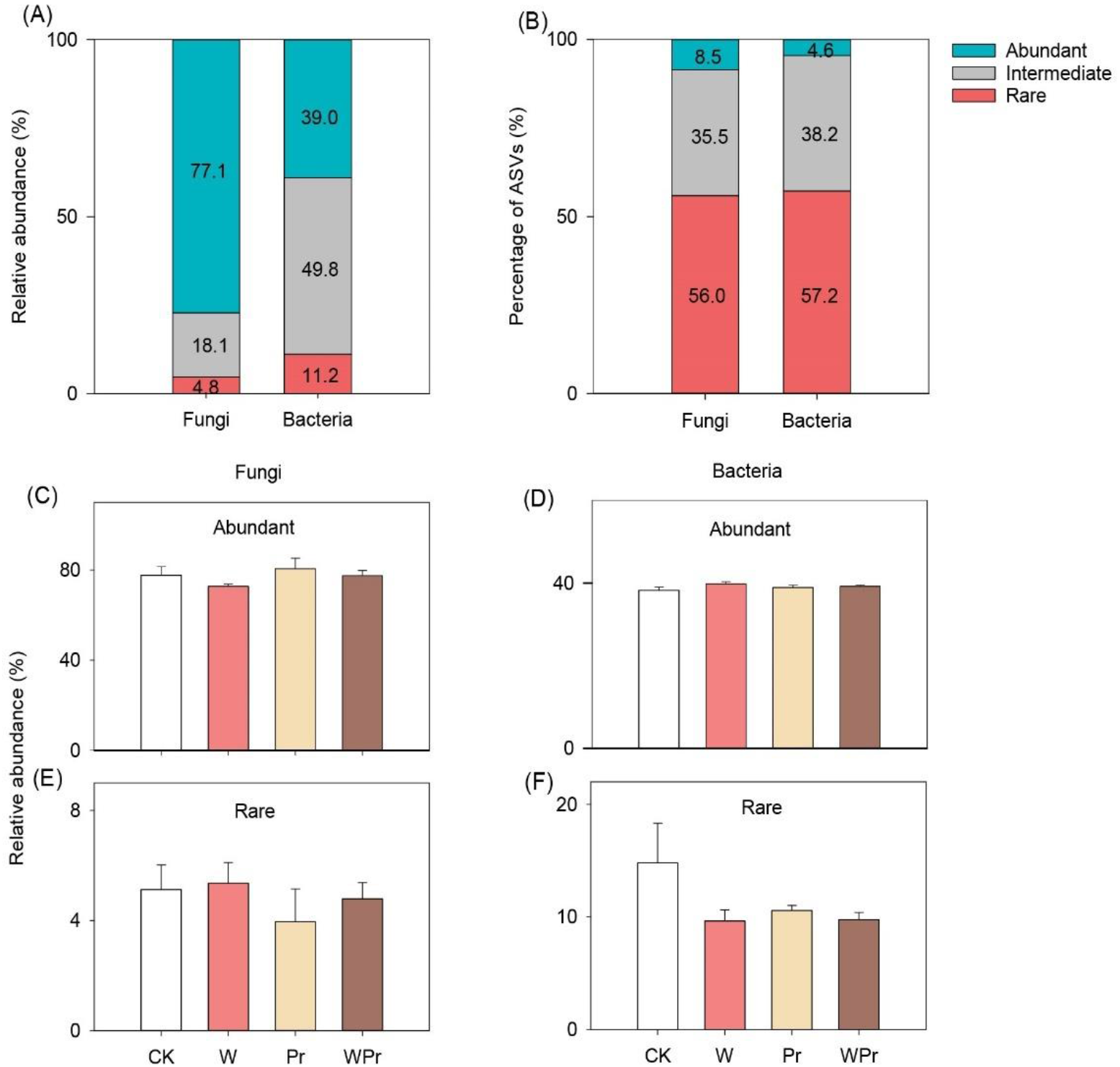
Relative abundance of abundant and rare taxa of soil fungi and bacteria. (A, B) The relative abundance and percentage of ASVs of abundant and rare taxa across the four treatments. (C-F) Effects of warming and precipitation reduction on the relative abundance of abundant and rare taxa of soil fungi and bacteria. Bars are standard errors of the means (n = 4). The four treatments are as follows: CK, control; W, warming; Pr, precipitation reduction; WPr, combination of warming and precipitation reduction.

Warming tended to reduce the abundance of abundant fungi by 5.1% (Fig. 3C), and precipitation reduction decreased the abundance of rare fungi by 16.6% (Fig. 3E). Neither warming nor precipitation reduction significantly impacted the abundance of abundant bacteria (Fig. 3D). Warming and precipitation tended to decrease the relative abundance of rare bacteria by 23.5% and 16.8%, respectively (Fig. 3F)

### Responses of alpha diversities of abundant and rare soil fungi and bacteria to warming and precipitation reduction

Across all the treatments, rare communities of fungi and bacteria had higher richness and Shannon indexes than abundant communities of fungi and bacteria, respectively (Figs. S2 and S3). Neither warming nor precipitation reduction significantly influenced the alpha diversity indices of abundant or rare fungi (i.e., richness, Shannon and Simpson indexes) (Fig. S2 and Table S4). While warming and precipitation reduction had no effect on alpha diversities of rare bacteria, warming significantly increased Simpson index and marginally increased Shannon index of abundant bacteria (Fig. S3 and Table S5).

### Responses of beta diversities of abundant and rare soil fungi and bacteria to warming and precipitation reduction

Warming and precipitation differently affected the beta diversities of fungi and bacteria. PERMANOVA showed that warming significantly altered abundant but not rare community composition of soil fungi, while precipitation reduction had effect on neither of them (Fig. 4A, 4C, 4E). PCoA revealed that abundant community composition of soil fungi was noticeably separated on axes 1 and 2, with 15.3% and 12.4% interpretations on axis 1 and 2, respectively (Fig. 4C). In addition, neither warming nor precipitation reduction had significant effects on abundant or rare community composition of soil bacteria (Fig. 4B, 4D, 4F).

**Fig. 4.**
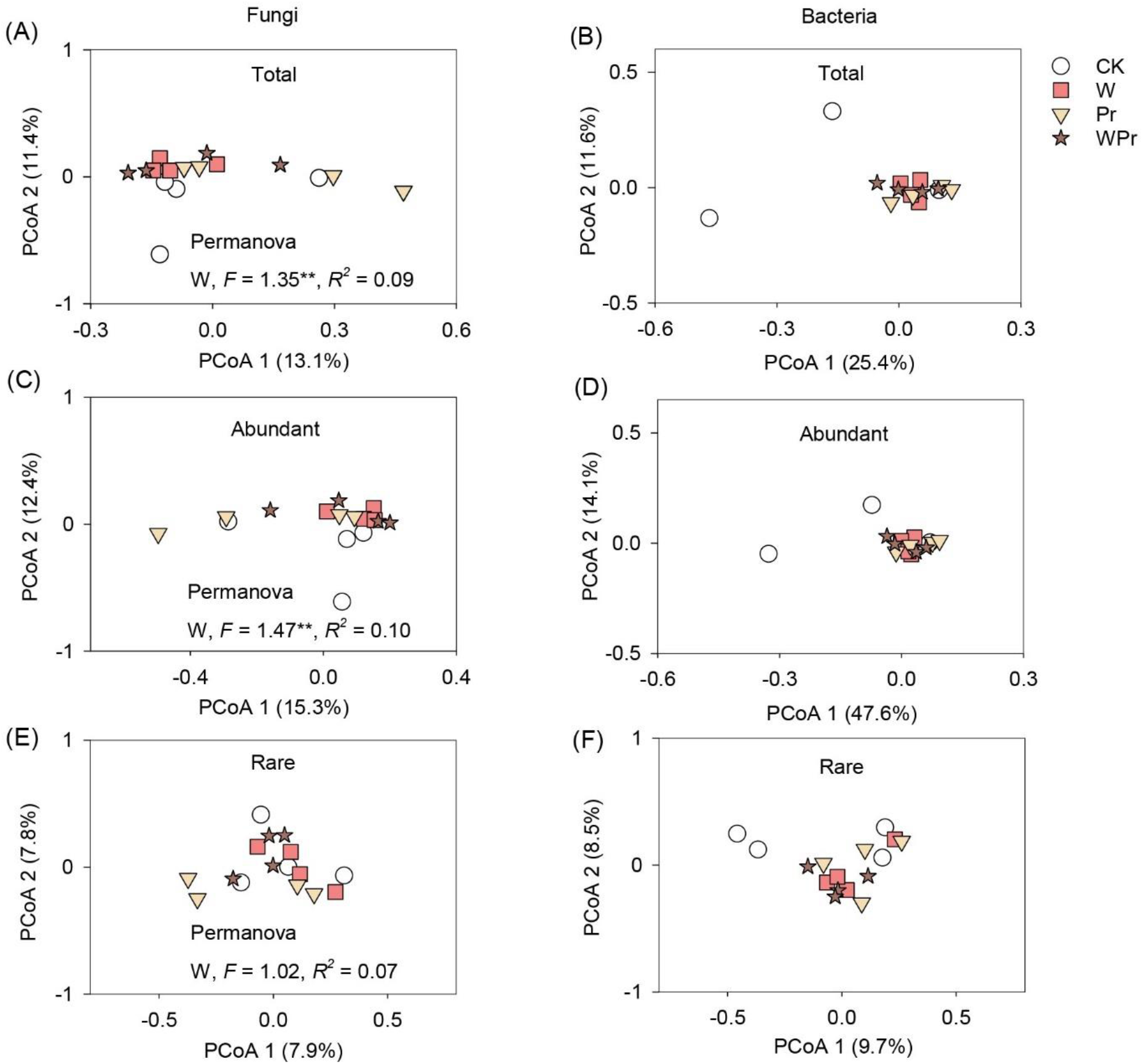
Soil fungal and bacterial community composition. Effects of warming and precipitation reduction on soil fungal and bacterial community composition (A, B: total; C, D: abundant; E, F: rare). Community composition of soil fungi and bacteria was assessed by PCoA at the taxon level based on the Bray–Curtis dissimilarity. The effects of warming and precipitation reduction on microbial composition were determined using PERMANOVA (*F*, *R*^2^, and *p* values; \*\**p* < 0.01). The four treatments are as follows: CK, control; W, warming; Pr, precipitation reduction; WPr, combination of warming and precipitation reduction.

In the fungal communities, *Ascomycota* (62.5%), *Mortierellomycota* (16.5%) and *Basidiomycota* (10.6%) were the most prevalent abundant fungal phyla (Fig. S4). The dominant rare fungal taxa were *Ascomycota* (46.1%), *Basidiomycota* (15.9%), *Mortierellomycota* (1.7%) and *Chytridiomycota* (1.5%) (Fig. S4). At the fungal order level, warming significantly increased the relative abundance of abundant *Capnodiales* and *Hypocreales*, but tended to decrease that of *Archaeorhizomycetales* (Fig. 5A-C and Table S6). Warming and precipitation reduction significantly increased the ratio of *Capnodiales* to *Archaeorhizomycetales* (Fig. 5D). Additionally, warming significantly increased the relative abundance of rare *Cantharellales* but decreased rare *Chaetothyriales* (Fig. S5 and Table S6).

**Fig. 5.**
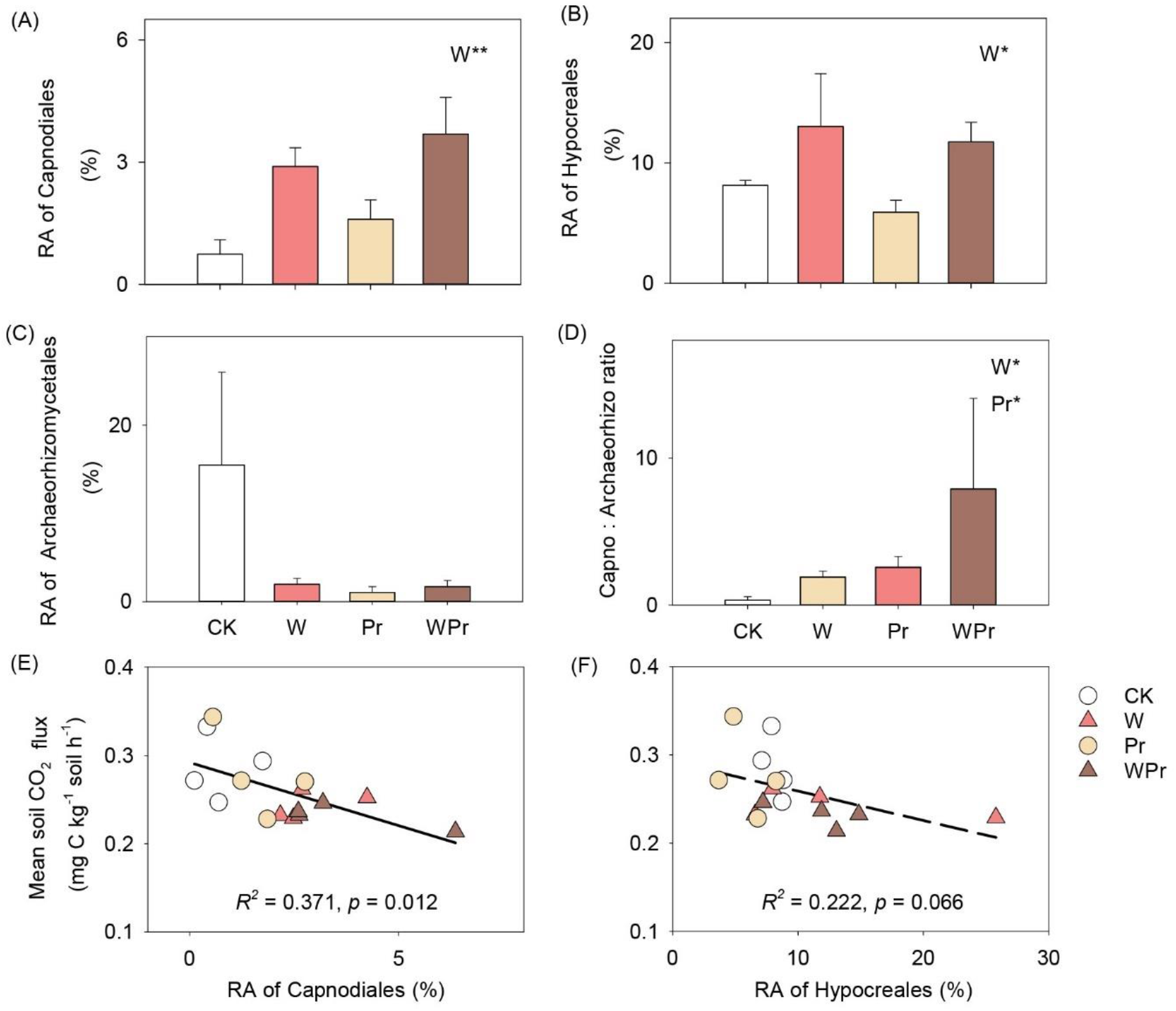
Relative abundance of domiant fungal *Capnodiales*, *Hypocreales*, *Archaeorhizomycetales*, and the ratio of *Capnodiales* to *Archaeorhizomycetales*. Effects of warming and precipitation reduction on the relative abundance of *Capnodiales* (A), *Hypocreales* (B), *Archaeorhizomycetales* (C), and the ratio of *Capnodiales* to *Archaeorhizomycetales* (D) in soil abundant fungal community, and the relationship between soil CO_2_ efflux and the relative abundance of *Capnodiales* (E) or *Hypocreales* (F), respectively. Bars are standard errors of the means (n = 4). The four treatments are as follows: CK, control; W, warming; Pr, precipitation reduction; WPr, combination of warming and precipitation reduction. The significance levels are labeled with *0.01 < *p* ≤ 0.05; **0.001 < *p* ≤ 0.01, ANOVA mixed model.

As for the bacterial communities, the rare community was dominated by phyla *Actinobacteria* (38.9%), *Proteobacteria* (23.2%), *Acidobacteria* (16.6%) and *Chloroflexi* (6.9%), which was similar to the abundant community with the four phyla accounting for 95.8% of the total sequences (Fig. S6). Neither warming nor precipitation reduction had any significant impacts on abundant or rare bacterial community composition at the phylum level (Fig. S6).

### Relationships among soil properties, fungal and bacterial abundance and community composition, and CO_2_ efflux

Our Pearson correlation analysis showed that soil CO_2_ efflux was significantly and positively correlated with soil available NH_4+_-N and the abundance of soil fungi, but negatively correlated with the relative abundance of soil abundant *Capnodiales* (*p* = 0.012) and *Hypocreales* (*p* = 0.066) (Fig. 5E, 5F). Mantel test revealed that abundant but not rare fungal community composition was correlated with soil CO_2_ efflux (Fig. 6 and Table S8). Also, the abundant fungal community composition was significantly correlated with soil available NH_4+_-N (Fig. 6 and Table S8). However, the rare fungal community composition showed a strong correlation with soil available NO_3-_-N (Fig. 6 and Table S8). Furthermore, our structural equation model (SEM) analysis showed that warming-induced decline in soil CO_2_ efflux was mainly explained by soil moisture, labile C and composition of abundant fungal community but not total fungal abundance (Fig. S7).

**Fig. 6.**
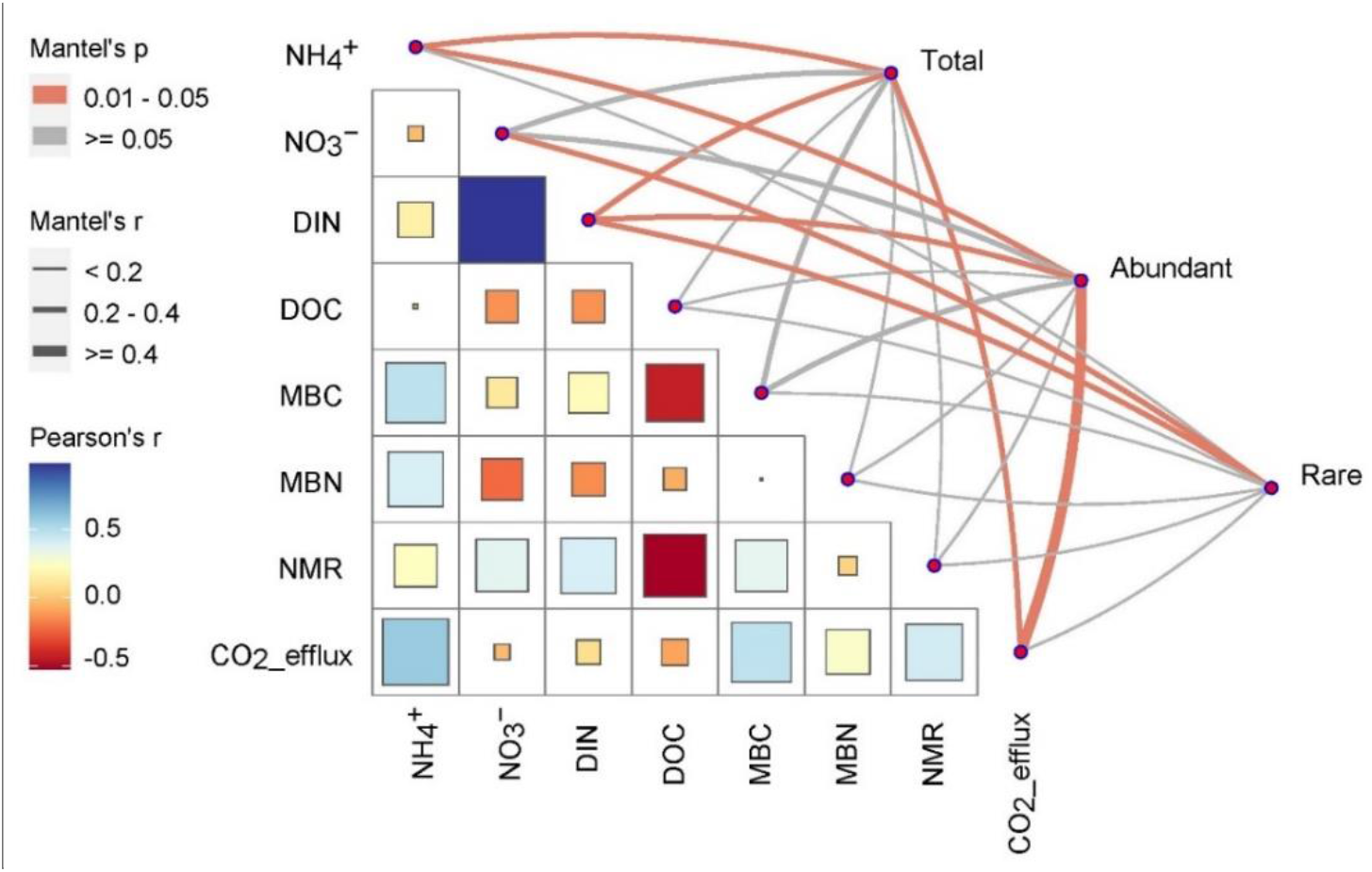
Potential drivers of fungal community composition. Relationships between fungal community composition (total, abundant and rare) and soil properties, root biomass and soil CO_2_ efflux. Correlations of soil fungal community structures (Bray-Curtis distance) with soil variables and ecosystem C fluxes. Edge width corresponds to the Mantel’s *r* value, and the edge colour denotes the statistical significance. Pairwise correlations of these variables are shown with a colour gradient presenting Pearson’s correlation coefficient. Soil variables include soil ammonium (NH_4+_), nitrate (NO_3-_), dissolved inorganic N (DIN), dissolved organic carbon (DOC), microbial biomass C (MBC) and N (MBN) and net mineralization rates (NMR).

## DISCUSSION

Microbial responses to climate warming and precipitation reduction may have the potential to significantly affect grassland ecosystem productivity and C dynamics. So far, studies have mainly focused on quantifying the alterations of soil microbial processes (e.g., soil respiration and N-cycling processes)^35,36^. Yet, the linkages among plant growth, soil microbes and soil C efflux, as influenced by climate warming and precipitation reduction, have not well been established.

Results from our field experiment showed that warming suppressed fungi, but not bacteria, and thus reduced soil C efflux in the semi-arid grassland (Figs. 1 and 2). These results were surprising and in contrast with our hypothesis because warming is expected to favor the growth of fungi over bacteria. Multiple mechanisms may have contributed to the observed shift in the microbial community structure (Figs. 2 and 4). Warming significantly increased root biomass at our site, while reducing soil moisture across the growing season^37^. This has likely increased root-derived organic C inputs to the soil as root exudates and fine root turnover increase, as evidenced by higher soil soluble C (Table 1). As bacteria and fungi are associated with different life strategies with bacteria growing and dividing faster than fungi, bacteria may outcompete fungi for more labile C utilization and/or via the production of antibiotics in the rhizosphere^38,39^. Also, evidence is mounting that antibiotics may be the primary driver shaping the soil microbial composition with a potential to affect ecosystem function^7,40,41^. Increased soil labile C availability and reduced moisture may prompt competition between bacteria and fungi for water and bacterial production of antibiotics may jointly play a major role in shaping the soil microbial community composition^42,43^. Further, according to the Preferential Substrate Utilization Hypothesis^44^, higher labile C and lower soil moisture under warming may promote microbes to utilize available labile C rather than to produce hydrolytic enzymes to decompose complex organic compounds, which may also contribute to the reduced CO_2_ efflux in our study.

Contrary to warming, precipitation reduction significantly increased soil MBC (Table 1) but had no effect on abundance, community composition of soil fungi and/or bacteria or soil C efflux (Figs. 1, 2 and 4). These results also contradict our expectation that reduced precipitation favors the growth of soil fungi over bacteria and suppress CO_2_ efflux (Hypothesis 1). These null effects of precipitation reduction on the microbial community indices may be due to a shift in plant community composition in favor of grasses over forbs under precipitation reduction at our site^36^. Compared with forbs, most grasses have highly branched fibrous roots, which are more fragile and turn over faster than taproots^36^. High root turnover likely increased plant C allocation belowground, contributing to the observed increase in soil MBC (Table 1). However, grasses have high efficiency in N uptake and may have induced relative N deficiency in microbes, as suggested by the higher MBC:MBN ratio (Table 1). Low N may further limit microbial production of enzymes and decomposition^4,7^, offsetting the positive effect on MBC. Together, our results suggest that while warming and precipitation reduction reduce soil moisture and constrain microbes, their indirect effects through affecting plant C allocation belowground and microbial interactions may play a more significant role in shaping the microbial community.

More interestingly, warming significantly altered the abundant fungal community composition (Fig. 4) and the abundant fungal community composition significantly correlated with soil CO_2_ emissions (by Mantel test, *p* < 0.05; Fig. 6). At this site, Bai et al. (2020)^37^ have previously observed that warming-driven reduction in total fungal biomass (estimated by the phospholipid fatty acids) correlated with decreased soil respiration. However, the relative effects on abundant and rare microbial taxa were not assessed. Different abundant and rare microbial taxa often vary in their sensitivity and strategies to soil temperature and moisture fluctuations^13,18^. Climate warming but not precipitation reduction resulted in preferable growth of order *Capnodiales* and *Hypocreales* over *Archaeorhizomycetales* (Fig. 5). *Capnodiales* (*Dothideomycetes*) and *Hypocreales* (*Sordariomycetes*) are considered as the potent degraders of recalcitrant C and important components of soil oligotrophic fungal taxa^45,46^. In a Tibetan alpine meadow, Che et al. (2019)^46^ reported that warming increased the proportion of an oligotrophic fungal class *Dothideomycetes* and reduced the proportion of active saprotrophic fungi. Compared with *Capnodiales* and *Hypocreales,* little is known about the ecology of the *Archaeorhizomycetales*, but it has been reported that some *Archaeorhizomycetales* may function as an important potential saprotroph in soil^47^. A decrease in the relative abundance of *Archaeorhizomycetales* under warming may partially contribute to the reduced soil CO_2_ emission (Figs. 1B and 5C). Additionally, it has been reported that *Capnodiales* are strong assimilators of plant residue C and may contribute positively to soil organic C sequestration^48^, similar to the negative correlation between soil CO_2_ efflux and the relative abundance of *Capnodiales* or *Hypocreales* observed in this study (Fig. 5E, 5F). Moreover, our SEM analysis confirmed that warming-induced decline in soil CO_2_ emissions was mainly due to alterations in soil C availability and moisture and abundant fungal community composition (Fig. S7).

In summary, our study represents one of the few field experiments that explicitly manipulated air temperature and reduced precipitation to examine the responses of different soil microbial subcommunities and CO_2_ efflux in semi-arid grasslands. Results from our field study illustrate that warming altered soil microbial communities, favoring the growth of bacteria over fungi. Also, warming induced a shift in the soil abundant fungal community composition in favor of *Capnodiales* and *Hypocreales* over potential saprotrophic *Archaeorhizomycetales*, which likely contributed to the reduced soil C efflux. Together, these results suggest that changes in soil C availability and moisture may dominate the microbial responses to climate change and thus control soil C dynamics in semi-arid grasslands under future climate change scenarios.

## MATERIALS AND METHODS

### The study site and the field experimental design

Our field study was carried out in a semi-arid grassland of Loess Plateau at the Yunwu Mountains Natural Preserve (36°13’N-36°19’N, 106°24’-106°28’E), northeast of Guyuan City, Ningxia Hui Autonomous Region of China, at approximately 2,000 m a.s.l. This experimental site has a typical semi-arid climate with a mean annual rainfall of about 455 mm. Approximately 70% of rainfall was concentrated in the growing season from July to September. The mean annual air temperature is 7 °C (ranging from -6.5 °C in January to 22.8 °C in July). The dominant plant species in the semi-arid grassland are *Artemisia sacrorum, Stipa przewalskyi, Stipa grandis, Chrysanthemus lavandulifolium and Saussurea alata*^49^. The study area has been fenced to exclude large animal grazing and protected since 1982. The soil at the study site is classified as a montane grey-cinnamon, Calci-Orthic Aridisol or a Haplic Calcisol based on Chinese and FAO classification system. The soil contained 40.2 g total C kg^-1^, 4.0 g total N kg^-1^, and had a pH of ca. 8.0 (measured in H_2_O).

The field experiment was set up on a largely flat hilltop (Fig. S1) in early June 2015 as a factorial experiment in a randomized block design with four blocks (replicates)^36,37,50^. Each block contains twelve treatment plots of 4 m × 4 m, separated by a buffer strip 1.5 m wide. There was also a 5 m buffer zone between each block. All treatments were randomly assigned to plots within blocks, and consisted of factorial combinations of two temperatures (ambient temperature and elevated temperature), three precipitation levels (precipitation addition (+30%), ambient precipitation and precipitation reduction (-30%)), and two N inputs (control and 12.0 N m^-2^ yr^-1^). Each warming treatment was achieved via open top chambers (OTCs) because electricity is inaccessible in this very remote field site. A large number of field studies have shown that this type of warming method was effective^37,50,51,52^. The OTCs were made of 6-mm-thick transparent polycarbonate materials with high light transmittance (>90%) and no infrared transmittance (<5%). They were hexagonal in shape, and had a height of 51.76 cm with 60 cm width at the top and 75 cm at the bottom (Fig. S1). There are four OTCs placed in each plot to ensure sufficient warming (Fig. S1). For the precipitation reduction (Pr) treatment, we placed rainout shelters made of seven tilted v-shaped transparent plexiglass 1 m above the soil surface on an iron hanger over each field Pr plot (see Fig. S1 for detail). Our previous publication showed that the rainout shelters intercepted approximately 30% of rainfall considering the wind effects on rain drops and possible lateral transport of water in field^36^. The rainwater was collected from each Pr plot using plastic containers and manually added to the nearest precipitation addition plot within 24-48 h after the rainfall event ended. Details of field N treatment have been described in previous publications^37,53^. For this study, only four treatments were selected for sampling: (a) ambient temperature and no precipitation reduction (CK); (b) warming (W); (c) precipitation reduction (Pr); (d) the combination of warming and precipitation reduction (WPr). Due to Covid-19 pandemic in 2020 that limited students’ access to the field plots, air, soil temperatures and field CO_2_ fluxes were not monitored. However, our previous record showed that air temperature increased by 2.0, 2.6 and 2.1°C, and reduced average soil volumetric moisture by 15.1, 16.8 and 13.5% in 2018, 2019 and 2021, respectively (Qiu et al. unpublished). It did not consistently increase averaged soil temperature at 10 cm depth in either 2018 or 2019, but significantly increased it by 0.5°C in 2021 (Qiu et al. unpublished). Also, our previous publication^36^ showed that precipitation reduction reduced soil moisture by 9.8, 10.9 and 7.6% in 2019, 2021 and 2022, respectively.

### Plant and soil sampling

In mid-August 2020 at the peak of the growing season, soil samples were collected from the field plots. In each plot, three 5-cm diameter soil cores (0-10 cm depth) were collected, and well mixed to generate a composite sample per plot. All the composite samples were immediately kept in coolers with ice bags and transported to the laboratory for processing. All samples were sieved to pass through a 2 mm sieve, and any visible stones and plant residues were carefully removed from the sieved soil. A small subsample (approximately 50 g) of soil was immediately frozen at -20 °C for molecular analysis. The remaining samples were then stored at 4 °C and used for the chemical and microbial analyses, which were conducted or initialized within 10 days.

### Soil physical, chemical and microbial analyses

Soil inorganic-N (NH_4+_-N and NO_3-_-N) and dissolved organic C (DOC) were determined from 20 g soils (dry soil equivalent), extracted with 50 mL of K_2_SO_4_ (0.5 M) after 30 min of rotary shaking. Soluble inorganic N and organic C in the extracts was quantified by a flow injection auto analyzer (SEAL-AA3, Germany) and a TOC analyzer (Elementar Vario TOC Cube), respectively. Soil microbial biomass C (MBC) and N (MBN) were determined using the chloroform fumigation extraction method^54^. Soil net N mineralization was assessed by the accumulation of inorganic N (NH_4+_-N and NO_3-_-N) during a 4-week incubation (25°C in the dark) with 20.0 g fresh soil following the method in Zhang et al. (2023)^36^.

### Soil CO_2_ efflux

Notes that field CO_2_ efflux in 2020 was not determined because the COVID limited students’ field access. Therefore, in this study, soil CO_2_ efflux rate was measured by the aerobic incubation method^55^. In brief, for each soil sample, equivalent of 20.0 g dried soil was weighed into a 120-ml mason jar and adjusted to a moisture level of 60% water holding capacity. All jars were then incubated at room temperature of 25 °C in the dark for 4 weeks. Soil CO_2_ emissions were measured after 1, 2, 4, 7, 10, 19 and 28 days of incubation following a static procedure as described in Zhang et al. (2023)^36^. CO_2_ concentrations were determined on a gas chromatograph (GC; 7890B; Agilent Technologies) equipped with a flame ionization detector (FID).

### Soil fungal and bacterial abundance

For each soil sample, total genomic DNA was extracted from 0.25 g (dry weight) frozen soil using the MoBio Power Soil DNA kit (MoBio Laboratories Inc, Carlsbad, USA) following the manufacturer’s instructions. Soil DNA quality and size were checked by electrophoresis on a 1% agarose gel. The abundance of total fungi and bacteria was determined using real-time quantitative PCR (qPCR) (7300 Real-Time PCR System; Applied Biosystems) of amplifying the fungal internal transcribed spacer (ITS) and bacterial 16S rRNA genes, respectively. Primer sets and PCR conditions for each gene are provided in Table S1. The PCR reactions were performed in triplicate and consisted of 12.5 µl of SYBRs Premix Ex Taq™ (Takara), 0.5 µl of Rox Reference Dye, 10.0 µl of ddH_2_O, 0.5 µl of each primer, and 1 µl of the template DNA, giving a final volume of 25 µl. Standard curves for qPCR were made following the method in Zhang et al. (2023)^36^. All PCR reaction efficiencies ranged between 95% and 110% with R^2^ > 0.98.

### Illumina MiSeq sequencing and analyses of fungal and bacterial communities

To characterize soil fungal and bacterial communities through high-throughput sequencing analysis, the fungal ITS and bacterial 16S rRNA genes were amplified using the same primer sets as for qPCR (see Table S1). All PCR reactions were carried out in triplicate with a total volume of 25 μL containing 5 μl of reaction buffer (5×), 5 μl of GC buffer (5×), 0.25 μl of Q5 DNA Polymerase, 2 μl of dNTPs (2.5 mM), 8.75 μl of ddH_2_O, 1 μl (10 μM) of each primer, and 2 μL of template DNA. The PCR amplification conditions were run with a 5 min initial denaturation at 98 °C, 25 cycles of denaturation at 98 °C for 30 s, annealing at 53 °C for 30 s, and extension at 72 °C for 45 s, and a final 5 min extension at 72 °C. The triplicate PCR products for each sample were combined and purified with Vazyme VAHTSTM DNA Clean Beads (Vazyme, Nanjing). The purified PCR products were then pooled in equimolar and sent for paired-end sequencing on the Illumina NovaSeq platform with NovaSeq 6000 SP Reagent Kit (Illumina, San Diego, CA) at Shanghai Personal Biotechnology Co., Ltd.

Raw sequencing data were processed using the Quantitative Insights Into Microbial Ecology 2 (QIIME2) version 2019.4.0^56^. In brief, raw sequences were demultiplexed using the demux plugin according to the unique barcodes for each sample. Then, the primers were cut from the sequences by the CUTADAPT plugin^57^. Sequences were quality filtered, denoised, merged, and chimera were removed using the DADA2 plugin^58^. And sequences below the quality score of 25 and fewer than 200 bp in length or sequences with ambiguous nucleotides were removed. After denoising, the filtered sequences were clustered into amplicon sequence variants (ASVs). The UNITE (http://unite.ut.ee/index.php)^59^ and SILVA databases (https://www.arb-silva.de/)^60^ were used to for fungal and bacterial taxonomy assignment.

The fungal and bacterial ASV tables were rarefied to 51,171 and 48,367 per sample for subsequent analyses, respectively. The abundant and rare ASVs were classified into six categories following recent publications^61–63^. Briefly, ASVs with relative abundances below 0.01% of the total sequences in all samples were designated as “always rare taxa, ART” and those with relative abundances < 0.01% in some samples but never ≥1% in any sample were “conditionally rare taxa, CRT”. ART and CRT were collectively referred to as rare taxa. ASVs with relative abundances ≥0.1% in all samples were designated as “always abundant taxa, AAT” and those with relative abundances ≥0.01% in all samples and ≥0.1% in some samples were “conditionally abundant taxa, CAT”. AAT and CAT were collectively referred to as rare taxa. The remaining ASVs (0.01%–0.1%) were defined as “intermediate taxa”. Here, we mainly focused on the abundant and rare taxa.

### Statistical analysis

Data were subjected to analysis of variance (ANOVA) using the stats package in R (version 4.1.2, R Development Core Team)^64^. Data were first tested for homogeneity of variances and normal distribution. Some variables were transformed for normal distribution where necessary. Plant root biomass and soil parameters were analyzed using linear mixed-effects model with warming and precipitation reduction treatments as the fixed factor and block as a random effect. For all tests, the significant differences were determined at the 95% probability level.

We analyzed changes in the composition and structure of fungal and bacterial communities between treatments, multivariate permutational analyses of the variance (PERMANOVA, 9,999 permutations) using the “adonis” function in the vegan package of R^65^. To visualize the differences in soil fungal and bacterial communities between experimental treatments, the principal coordinate analysis (PCoA) using Bray-Curtis distances was employed. For this, we used the “metaMDS” function in the vegan package^65^. Relationships between soil fungal communities and soil variables were examined by Mantel test with “mantel” function in the vegan package^65^.

Structural equation models (SEMs) analyses were further conducted to examine the effects of warming on soil CO_2_ emission via biotic and abiotic variables. We used PCoA axis 1 scores as a proxy for abundant fungal community composition. The data were fitted to the models using the maximum likelihood estimation method and fitted with the χ^2^ test. The SEM analysis was performed with the lavaan package^66^ in R.

## ACKNOWLEDGMENTS

We thank Qilai Yang, Hongdong Ma, Luyao Zhou, Hui Guo, Zhen Li and Fuwei Wang for their assistance in field experimental maintenance. We are thankful to the Ningxia Yunwu Mountains Grassland Natural Reserve Administration for logistical support. This work was supported by and National Natural Science Foundation of China (NSFC) (No.32371626, 32001140) and China Postdoctoral Science Foundation (No. 2022T150325).

## AUTHOR CONTRIBUTIONS

S.H. and Y.W. established the long-term field experiment. Y.Q. and S.H. conceived and designed the study. Y.Q., K.Z., Yunfeng Zhao, Yexin Zhao, B.W., T.H., X.X. and T.B. performed field and lab work. Y.Q., K.Z. and Yexin Zhao conducted the bioinformatic and statistical analysis. Y.Q. and S.H. wrote the manuscript with significant inputs from all other authors.

## ETHICS STATEMENT

This article does not contain any studies with human participants or animals performed by any of the authors.

## DATA AVAILABILITY

The representative sequences of ITS and 16S rRNA gene amplicons were available in figshare (https://doi.org/10.6084/m9.figshare.23257013).All other relevant data are available upon reasonable request from the corresponding author (Yunpeng Qiu or Shuijin Hu).

## SUPPORTING INFORMATION

Additional Supporting Information is available

Figs. S1-S7

Tables S1-S8

## ORCID

Yunpeng Qiu: https://orcid.org/0000-0003-2436-6579 Shuijin Hu: https://orcid.org/0000-0002-3225-5126

## Additional Supporting Information

**Figure S1.**
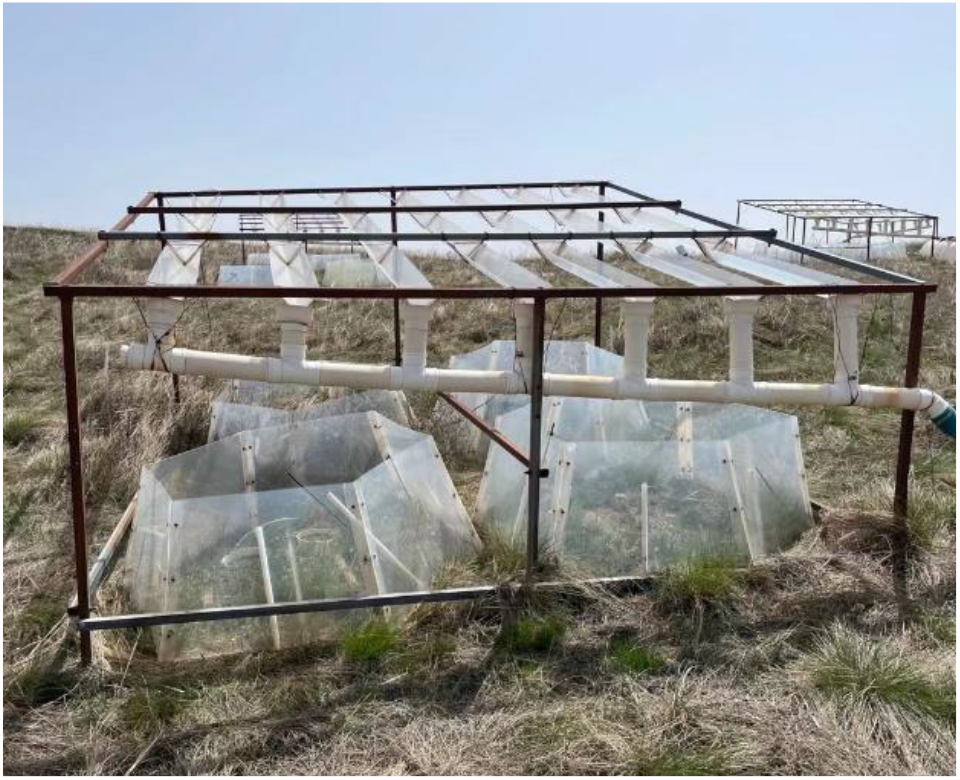
A partial overview of field warming and precipitation reduction plots. Warming was simulated via open-top chambers (OTCs). Seven tilted v-shaped transparent plexiglass were placed 1 m high above soil surface to intercept 30% of rainfall to simulate a light to moderate precipitation reduction. Photo credit: Kangcheng Zhang.

**Figure S2.**
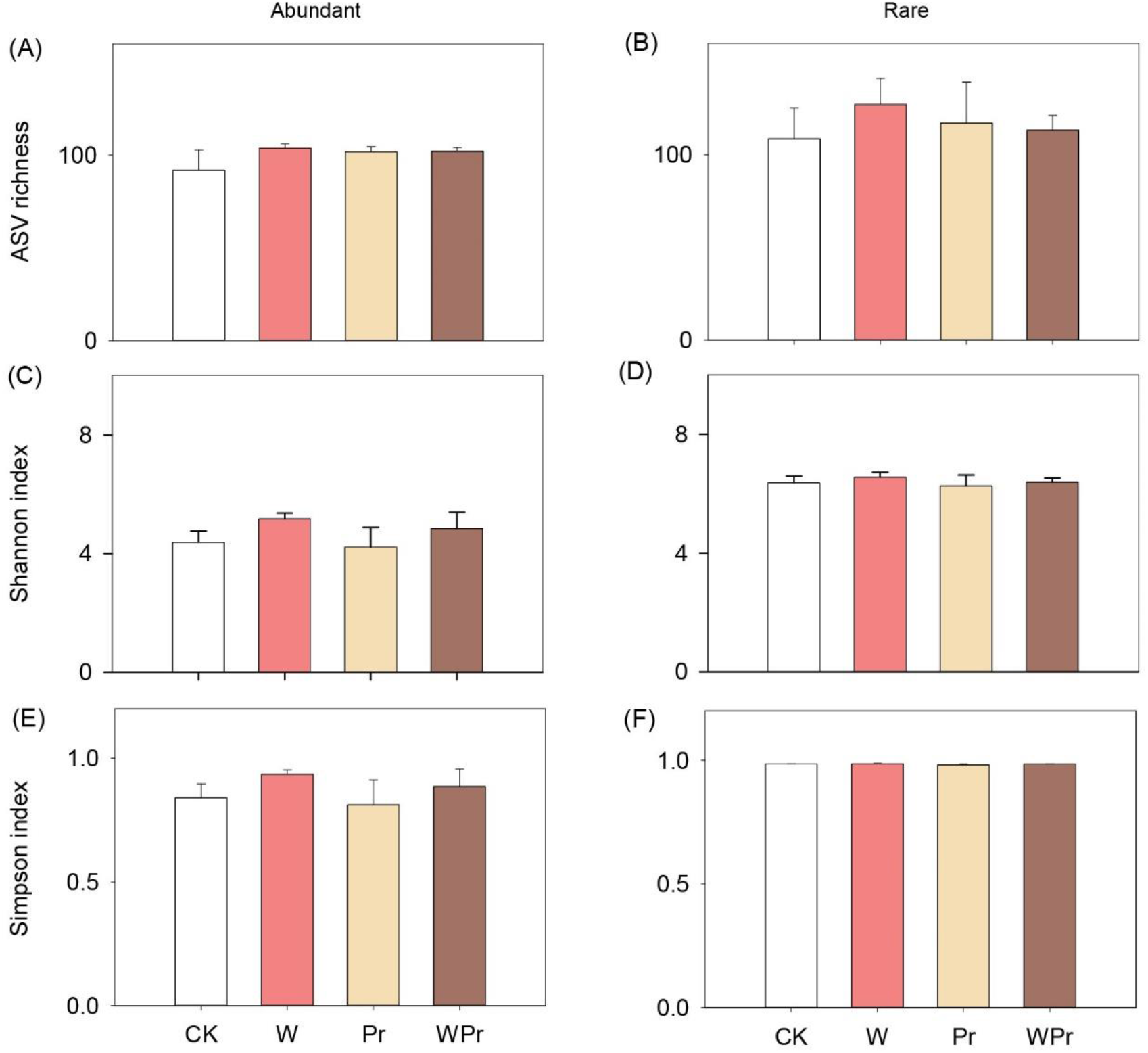
Soil fungal alpha diversity. Effects of warming and precipitation reduction on abundant and rare fungal alpha diversity. Values are means ± SE (n = 4). The four treatments are as follows: CK, control; W, warming; Pr, precipitation reduction; WPr, combination of warming and precipitation reduction.

**Figure S3.**
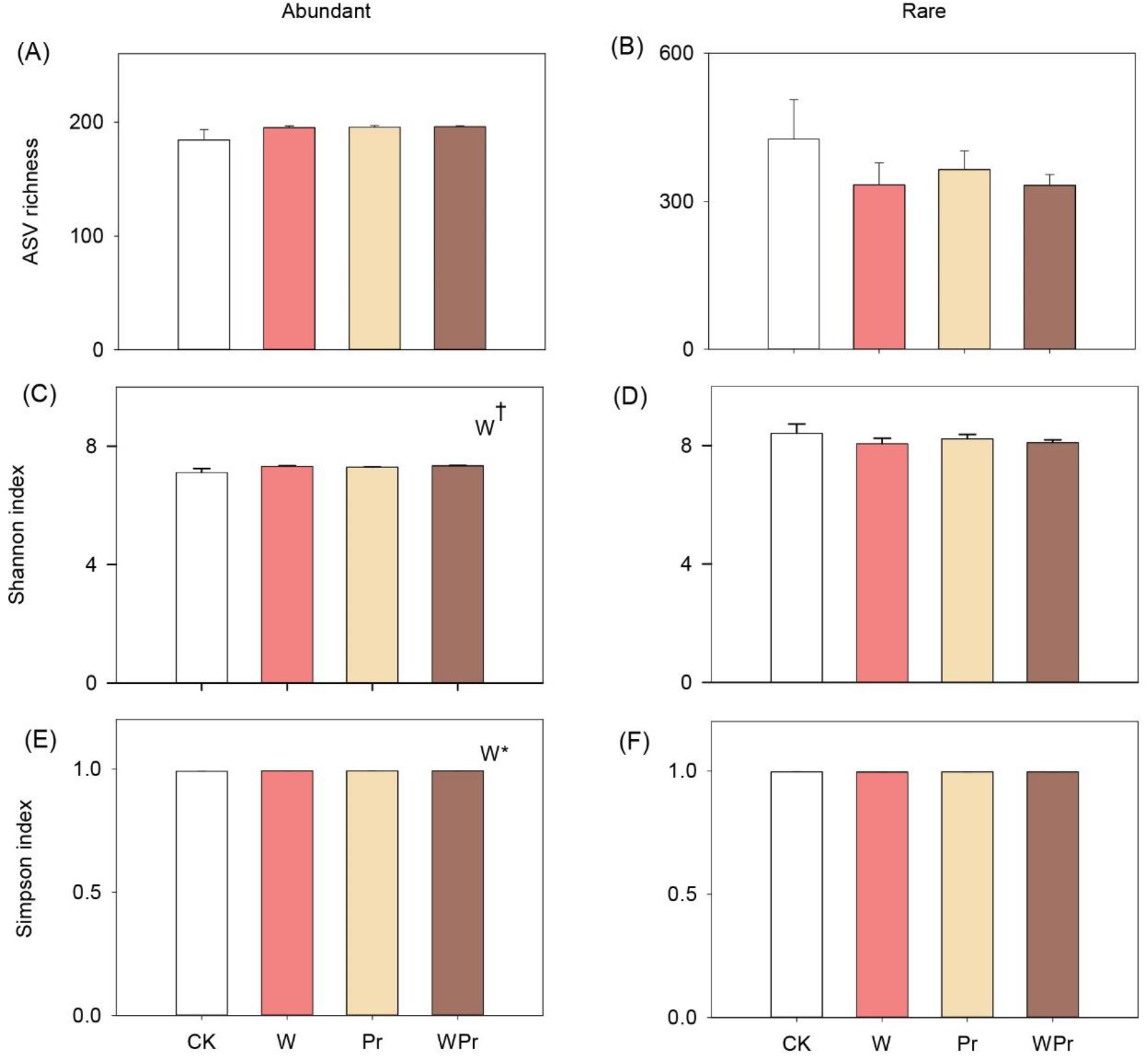
Soil bacterial alpha diversity. Effects of warming and precipitation reduction on abundant and rare bacterial alpha diversity. Values are means ± SE (n = 4). The four treatments are as follows: CK, control; W, warming; Pr, precipitation reduction; WPr, combination of warming and precipitation reduction. The significance levels are labeled with †0.05 < *p* ≤ 0.10; *0.01 < *p* ≤ 0.05, ANOVA mixed model.

**Figure S4.**
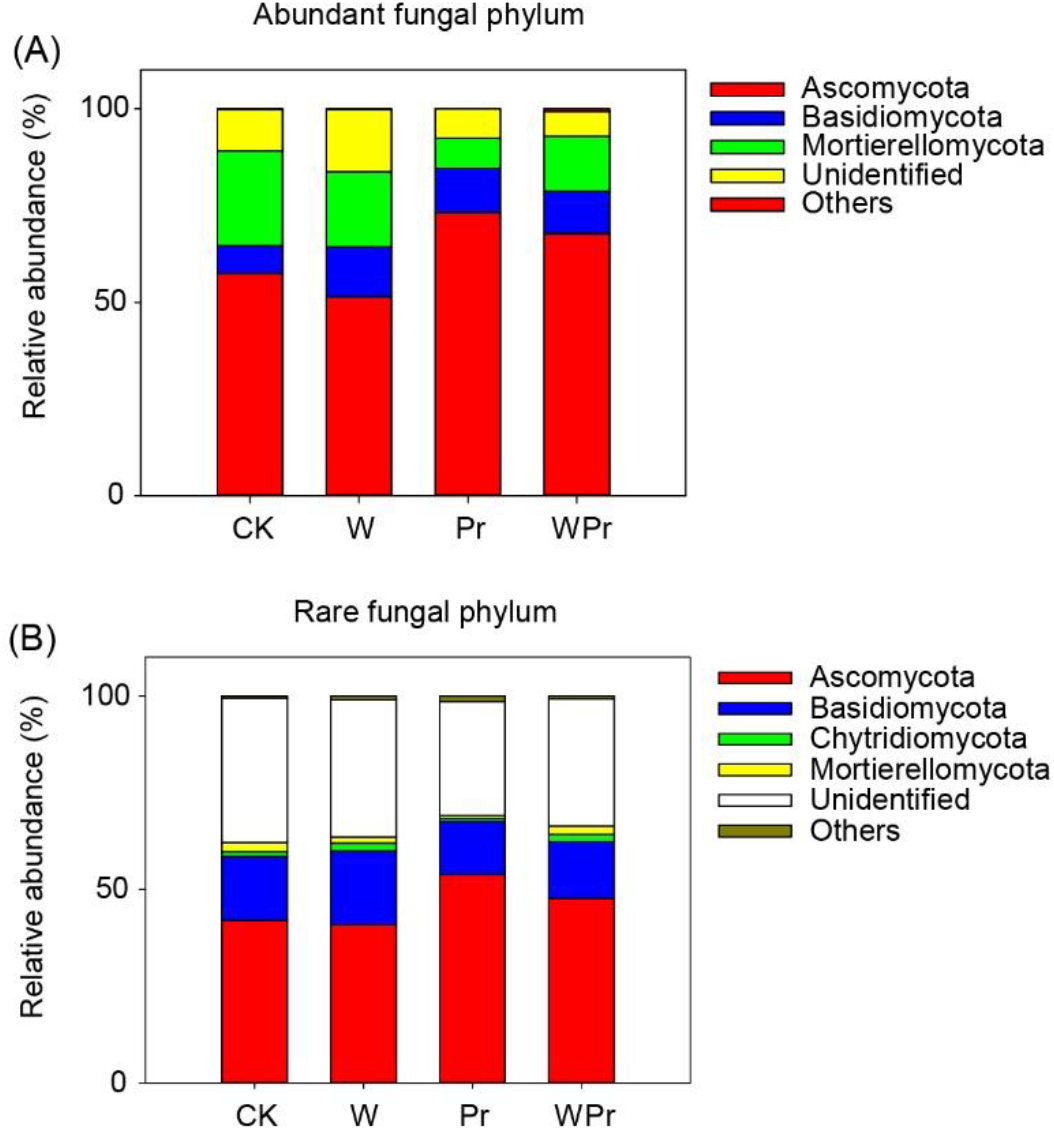
Soil fungal community composition at phylum level. Effects of warming and precipitation reduction on the composition of abundant and rare fungal taxa at phylum level. The four treatments are as follows: CK, control; W, warming; Pr, precipitation reduction; WPr, combination of warming and precipitation reduction.

**Figure S5.**
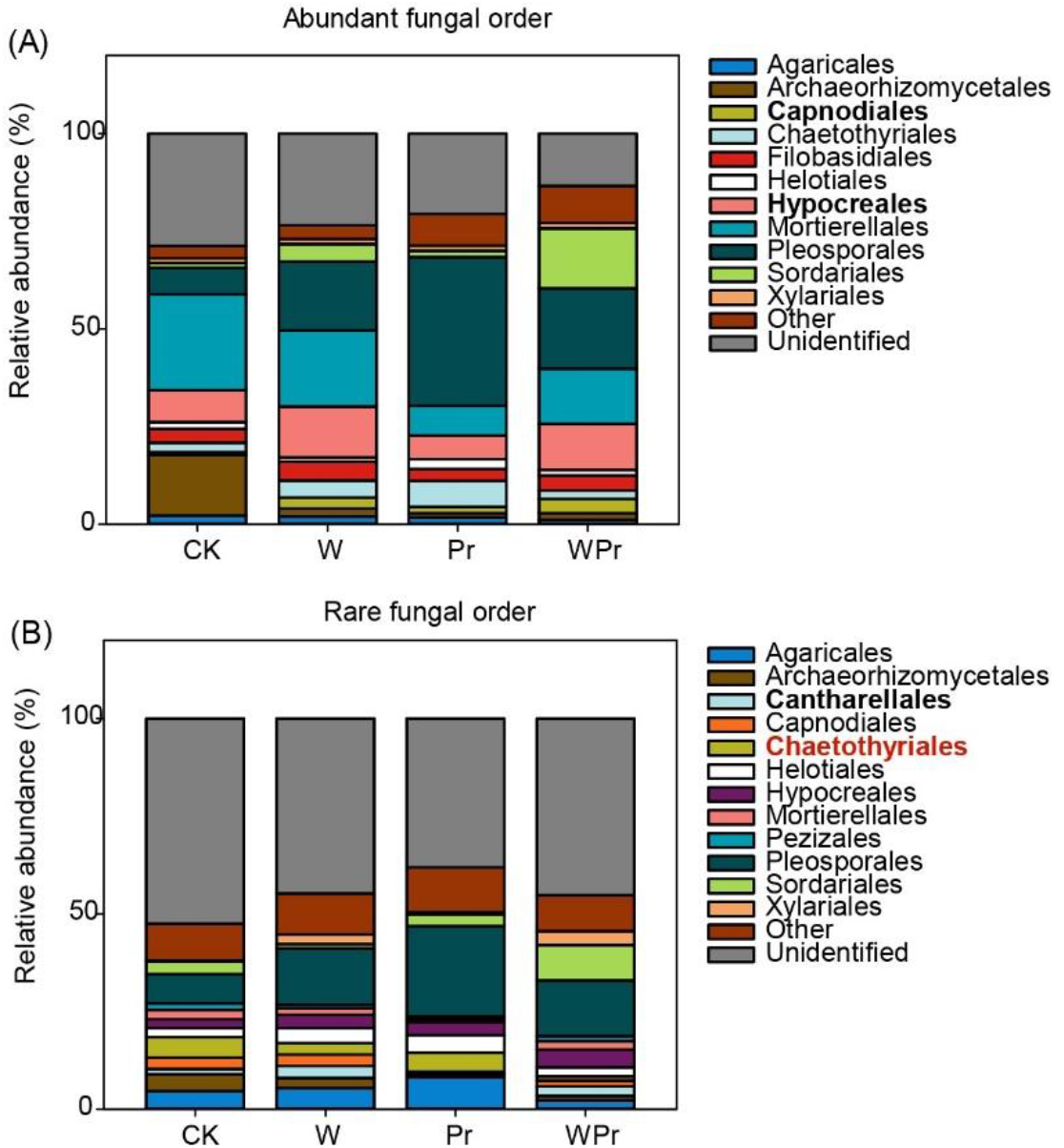
Soil fungal community composition at order level. Effects of warming and precipitation reduction on the composition of abundant (A) and rare (B) fungal taxa at order level. Black bold denotes positive effect of warming on the order and the red bold denotes negative effect of warming on the order. The four treatments are as follows: CK, control; W, warming; Pr, precipitation reduction; WPr, combination of warming and precipitation reduction.

**Figure S6.**
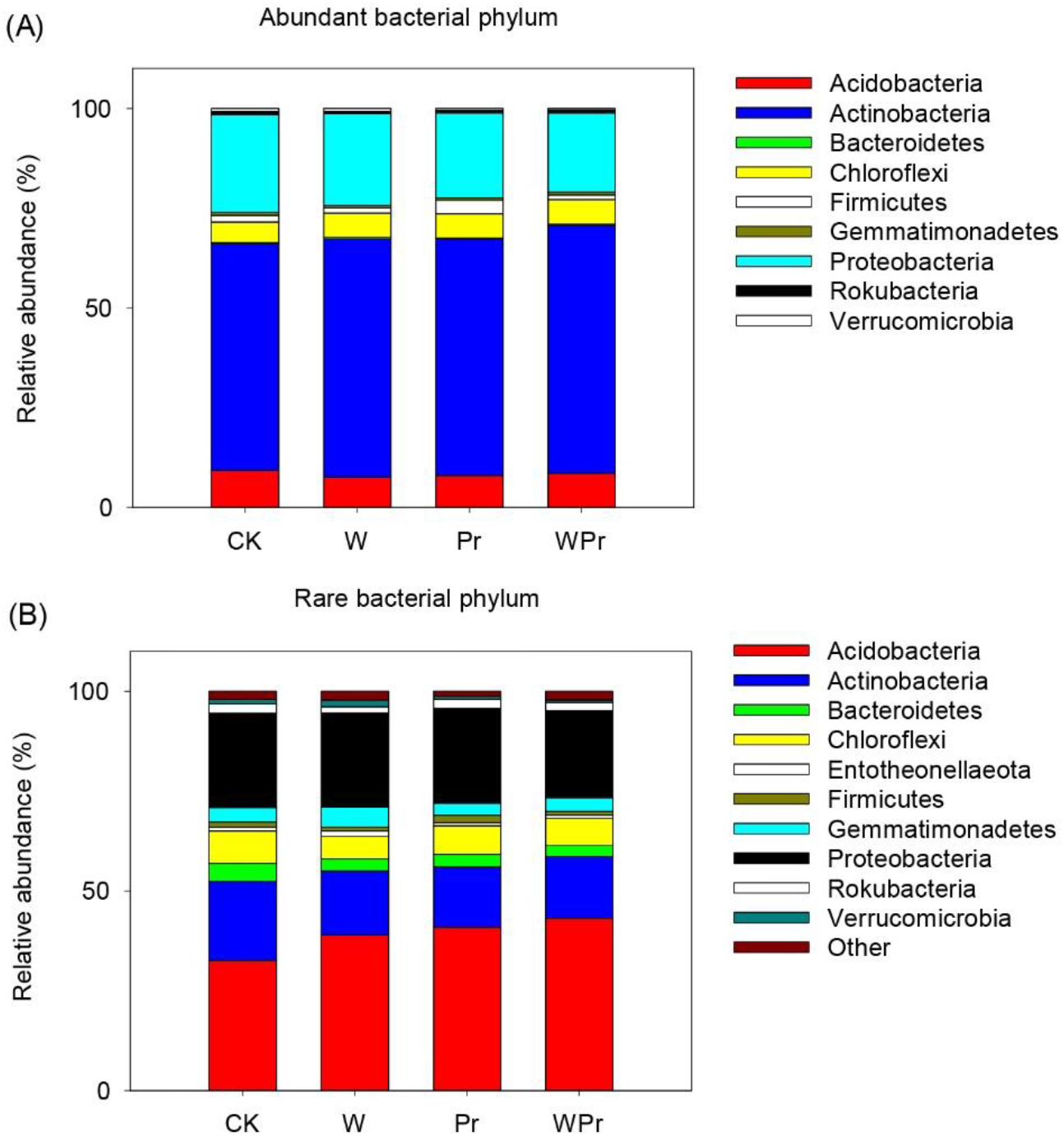
Soil bacterial community composition at phylum level. Effects of warming and precipitation reduction on the composition of abundant (A) and rare (B) bacterial taxa at phylum level. The four treatments are as follows: CK, control; W, warming; Pr, precipitation reduction; WPr, combination of warming and precipitation reduction.

**Figure S7.**
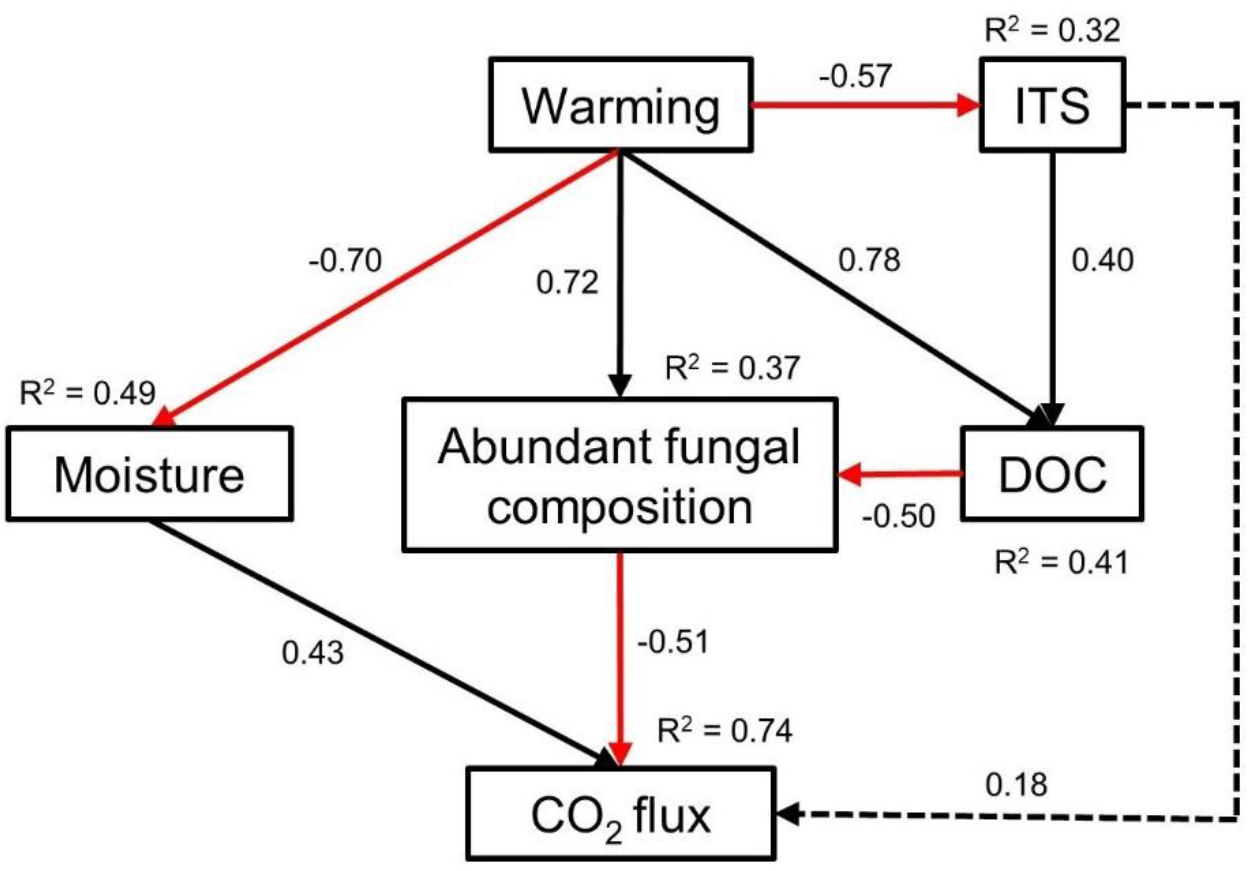
Potential drivers of soil CO_2_ flux. Structural equation modelling (SEM) analysis of the effects of warming on soil CO_2_ emissions. Results of the final model fitting: Chi-square (χ^2^) = 6.658, *P* = 0.354, degree of freedom (df) = 6, comparative fit index (CFI) = 0.986, root square mean error of approximation (RMSEA) = 0.083. Abundant fungal community composition is indicated by the first axis of PCoA. ITS, the abundance of soil fungi; DOC, dissolved organic carbon. Values associated with arrows are standardized path coefficients. Black arrows indicate significant positive relationships and red arrows indicate significant negative relationships (*P* < 0.05). The dashed arrows indicate non-significant relationship (*P* > 0.05). *R*^2^ values are the proportion of variation explained by relationships with other variables.

**Table S1.**
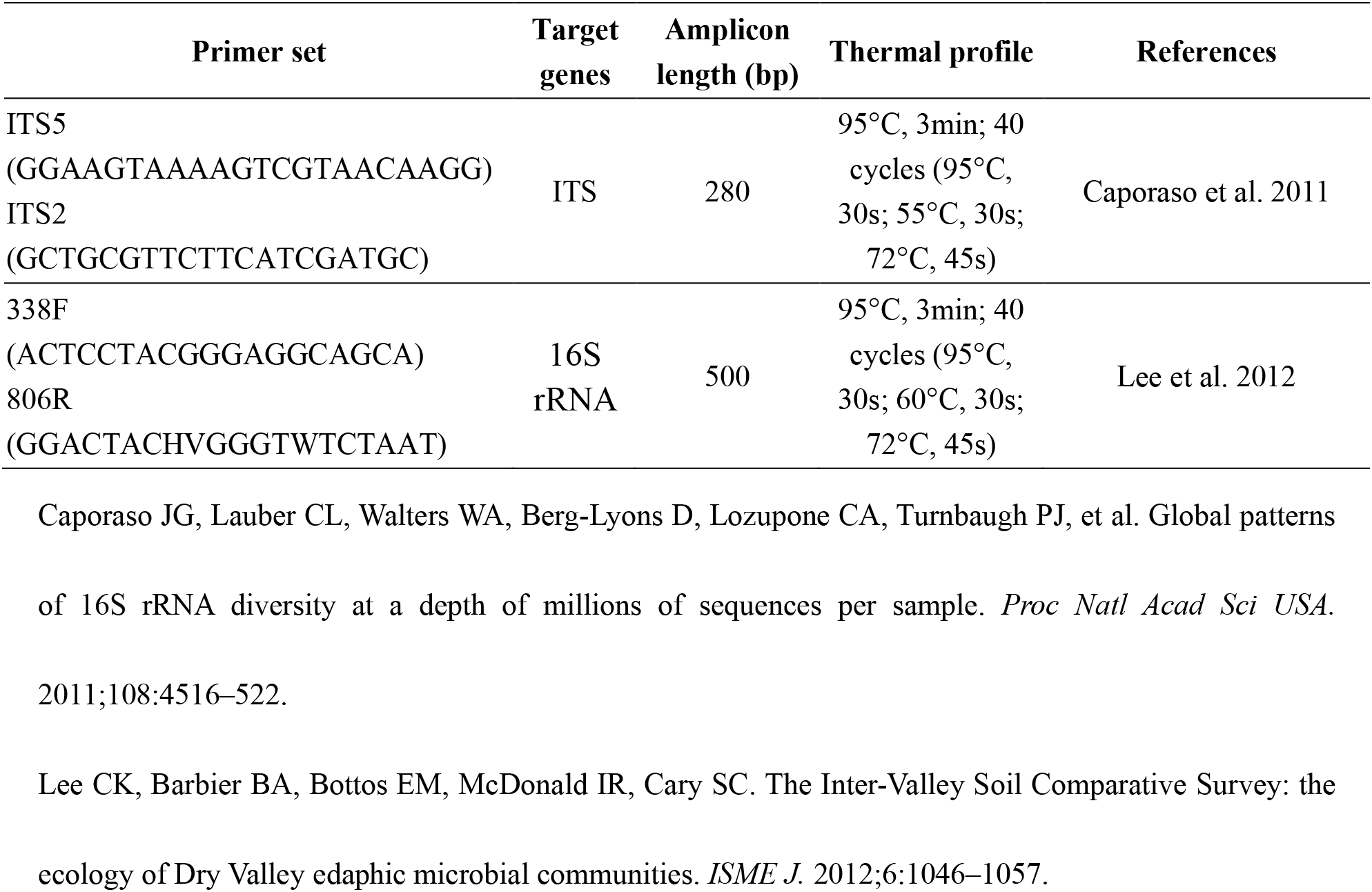
Primers and conditions used for real-time quantitative PCR of fungal ITS and genes.

**Table S2.** Results (*P* values) of ANOVA for the effects of warming and precipitation reduction (Pr) on the soil bacterial, fungal abundance, and the ratio of fungal to bacterial abundance, and mean soil CO_2_ flux.

| Variables | W | Pr | W × Pr |
| --- | --- | --- | --- |
| 16S rRNA | 0.735 | 0.975 | 0.230 |
| ITS | **0.009** | 0.576 | **0.027** |
| ITS/16S rRNA | **0.019** | 0.671 | 0.377 |
| Mean CO<sub>2</sub> flux | **0.020** | 0.485 | 0.824 |

**Table S3.** Results (*P* values) of ANOVA for the effects of warming and precipitation reduction (Pr) on the relative abundance of abundant and rare fungal and bacterial taxa.

|  | Variables | W | Pr | W × Pr |
| --- | --- | --- | --- | --- |
| Fungi | Abundant | 0.237 | 0.266 | 0.789 |
|  | Rare | 0.544 | 0.335 | 0.732 |
| Bacteria | Abundant | 0.138 | 0.970 | 0.307 |
|  | Rare | 0.124 | 0.275 | 0.247 |

**Table S4.**
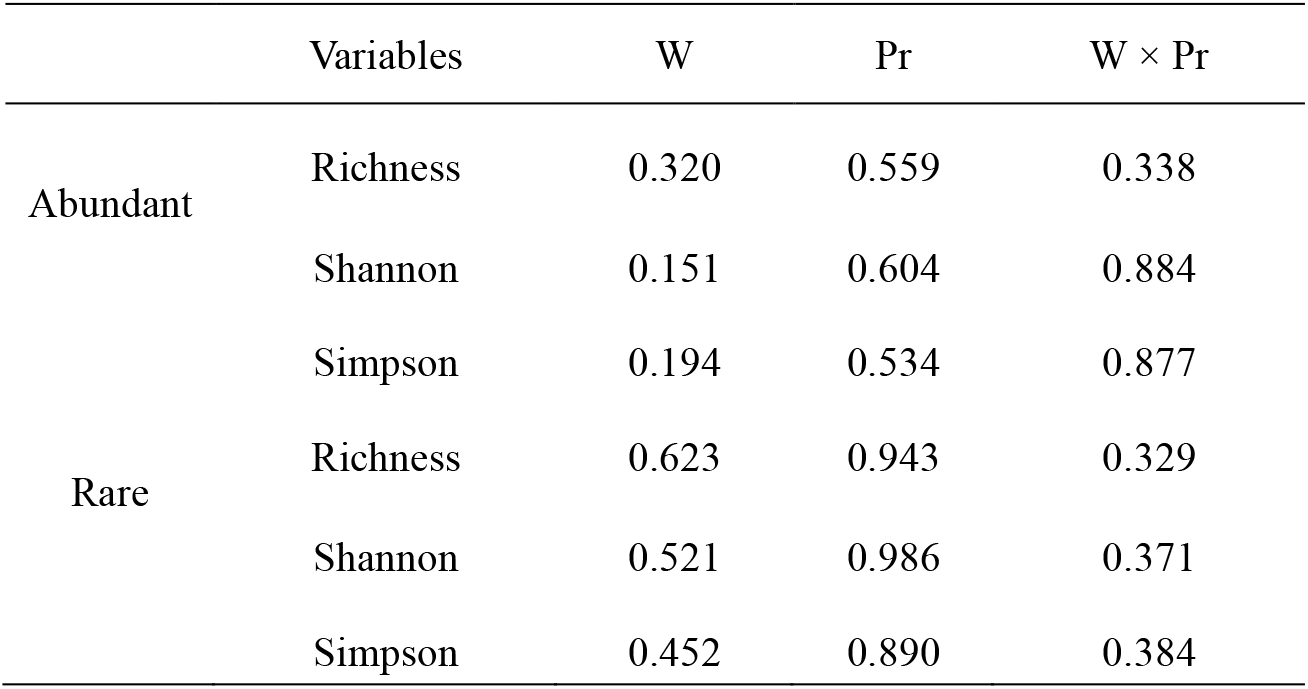
Results (*P* values) of ANOVA for the effects of warming and precipitation reduction (Pr) on alpha diversities of abundant and rare fungi.

|  | Variables | W | Pr | W × Pr |
| --- | --- | --- | --- | --- |
| Abundant | Richness | 0.320 | 0.559 | 0.338 |
|  | Shannon | 0.151 | 0.604 | 0.884 |
|  | Simpson | 0.194 | 0.534 | 0.877 |
| Rare | Richness | 0.623 | 0.943 | 0.329 |
|  | Shannon | 0.521 | 0.986 | 0.371 |
|  | Simpson | 0.452 | 0.890 | 0.384 |

**Table S5.**
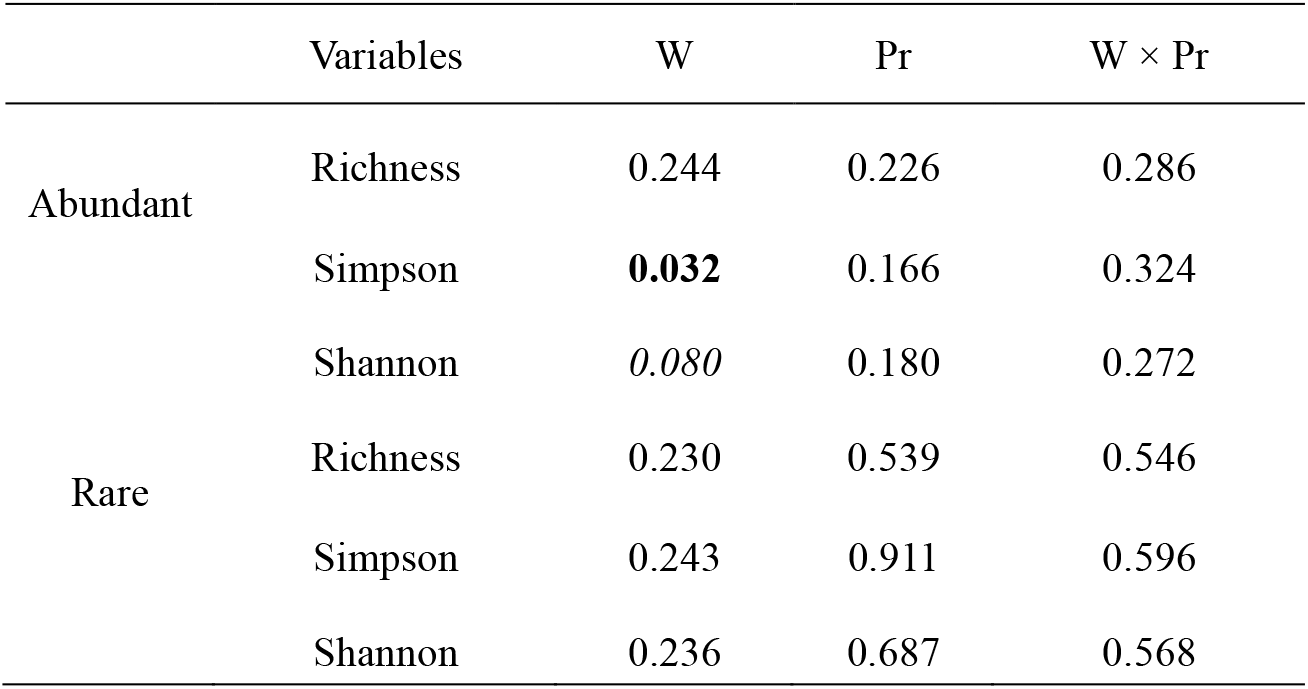
Results (*P* values) of ANOVA for the effects of warming and precipitation reduction (Pr) on alpha diversities of abundant and rare bacteria.

|  | Variables | W | Pr | W × Pr |
| --- | --- | --- | --- | --- |
| Abundant | Richness | 0.244 | 0.226 | 0.286 |
|  | Simpson | **0.032** | 0.166 | 0.324 |
|  | Shannon | *0.080* | 0.180 | 0.272 |
| Rare | Richness | 0.230 | 0.539 | 0.546 |
|  | Simpson | 0.243 | 0.911 | 0.596 |
|  | Shannon | 0.236 | 0.687 | 0.568 |

**Table S6.**
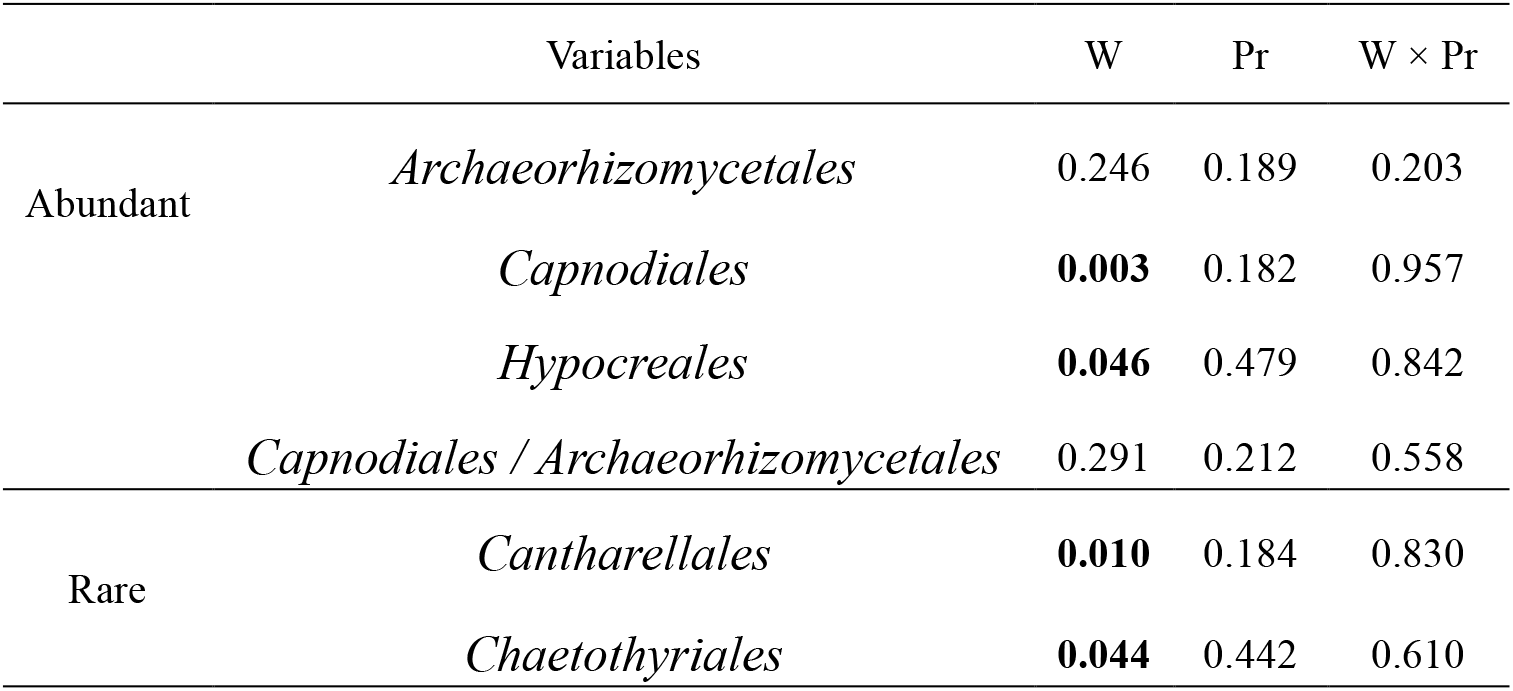
Results (*P* values) of ANOVA for the effects of warming and precipitation reduction (Pr) on soil abundant (i.e., *Archaeorhizomycetales, Capnodiales* and *Hypocreales*) and rare fungal order (i.e., *Cantharellales* and *Chaetothyriales*).

**Table S7.**
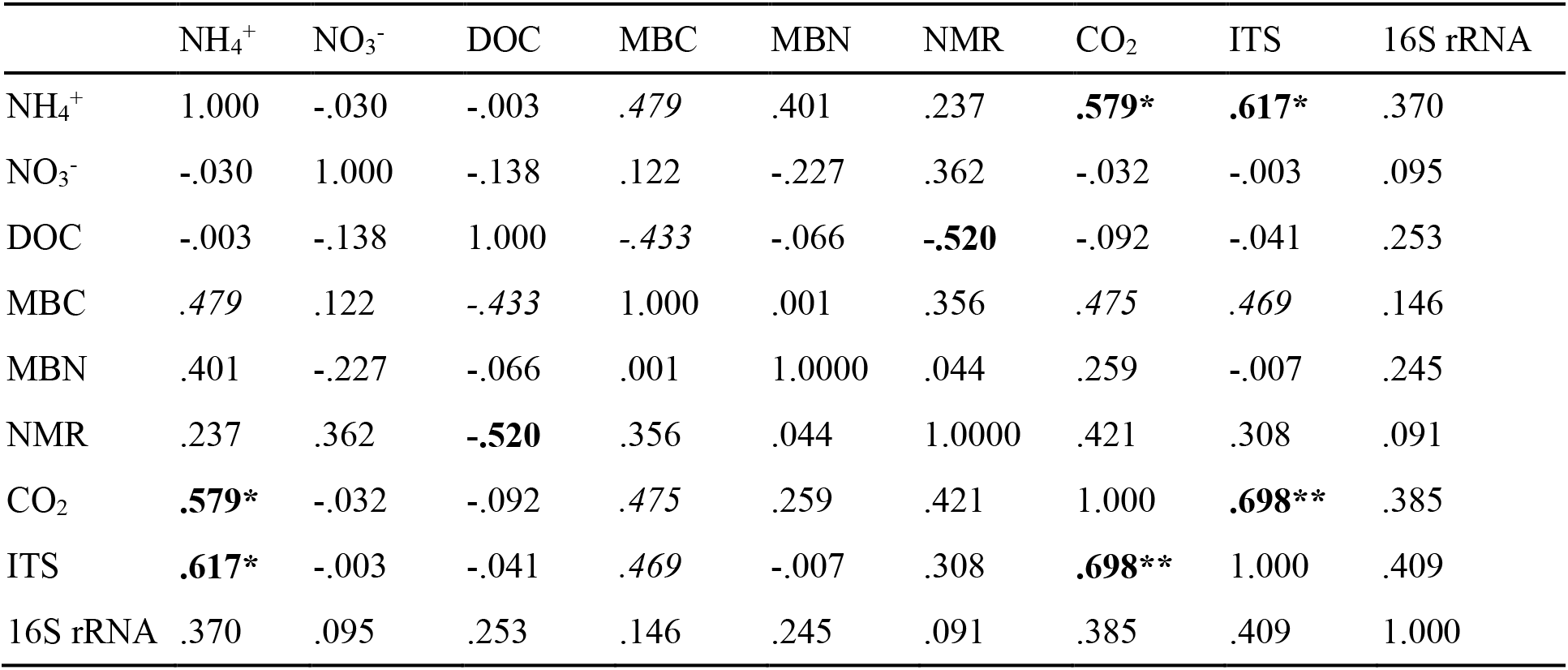
Pearson correlations among soil N pools, transformation and microbial parameters.

|  | NH<sub>4</sub><sup>+</sup> | NO<sub>3</sub><sup>-</sup> | DOC | MBC | MBN | NMR | CO<sub>2</sub> | ITS | 16S rRNA |
| --- | --- | --- | --- | --- | --- | --- | --- | --- | --- |
| NH<sub>4</sub><sup>+</sup> | 1.000 | -.030 | -.003 | .479 | .401 | .237 | **.579\*** | **.617\*** | .370 |
| NO<sub>3</sub><sup>-</sup> | -.030 | 1.000 | -.138 | .122 | -.227 | .362 | -.032 | -.003 | .095 |
| DOC | -.003 | -.138 | 1.000 | -.433 | -.066 | **-.520** | -.092 | -.041 | .253 |
| MBC | .479 | .122 | -.433 | 1.000 | .001 | .356 | .475 | .469 | .146 |
| MBN | .401 | -.227 | -.066 | .001 | 1.0000 | .044 | .259 | -.007 | .245 |
| NMR | .237 | .362 | **-.520** | .356 | .044 | 1.0000 | .421 | .308 | .091 |
| CO<sub>2</sub> | **.579\*** | -.032 | -.092 | .475 | .259 | .421 | 1.000 | **.698\*\*** | .385 |
| ITS | **.617\*** | -.003 | -.041 | .469 | -.007 | .308 | **.698\*\*** | 1.000 | .409 |
| 16S rRNA | .370 | .095 | .253 | .146 | .245 | .091 | .385 | .409 | 1.000 |

**Table S8.** Mantel test (Bray-Curtis) results for the relationships between fungal community composition (total, abundant and rare) and soil properties and soil CO_2_ efflux.

| Variables | Total |  | Abundant |  | Rare |  |
| --- | --- | --- | --- | --- | --- | --- |
|  | *r value* | *P value* | *r value* | *P value* | *r value* | *P value* |
| NH<sub>4</sub><sup>+</sup>-N | 0.312 | **0.049** | 0.331 | **0.034** | 0.016 | 0.425 |
| NO<sub>3</sub><sup>-</sup>-N | 0.308 | 0.076 | 0.299 | 0.082 | 0.311 | **0.047** |
| DIN | 0.368 | **0.045** | 0.358 | **0.049** | 0.329 | **0.043** |
| DOC | -0.003 | 0.477 | -0.010 | 0.486 | -0.047 | 0.623 |
| MBC | 0.225 | 0.131 | 0.240 | 0.114 | 0.125 | 0.170 |
| MBN | -0.038 | 0.555 | -0.033 | 0.540 | -0.063 | 0.680 |
| NMR | 0.113 | 0.275 | 0.117 | 0.258 | -0.045 | 0.592 |
| CO<sub>2</sub> efflux | 0.397 | **0.035** | 0.419 | **0.027** | 0.096 | 0.254 |

DIN, dissolved inorganic nitrogen; DOC, dissolved organic carbon; NMR, net nitrogen mineralization rates; MBN, soil microbial biomass nitrogen; MBC, soil microbial biomass carbon. Bold values indicate a significant different at $P < 0.05$ .

